# The integrity of dopaminergic and noradrenergic brain regions is associated with different aspects of late-life memory performance

**DOI:** 10.1101/2022.10.12.511748

**Authors:** Martin J. Dahl, Shelby L. Bachman, Shubir Dutt, Sandra Düzel, Nils C. Bodammer, Ulman Lindenberger, Simone Kühn, Markus Werkle-Bergner, Mara Mather

**Author notes:** Corresponding author: MJD.

## Abstract

Researchers have identified changes in dopaminergic neuromodulation as playing a key role in adult memory decline. Facilitated by technical advancements, recent research has also implicated noradrenergic neuromodulation in shaping late-life memory performance. However, it is not yet clear whether these two neuromodulators have distinct roles in age-related cognitive changes.

Combining longitudinal high-resolution magnetic resonance imaging of the dopaminergic substantia nigra–ventral tegmental area (SN–VTA) and the noradrenergic locus coeruleus (LC) in younger (n = 69) and older adults’ (n = 251), we found that dopaminergic and noradrenergic integrity are differentially associated with individual differences in memory performance. While LC integrity was related to better episodic memory across several memory tasks, SN–VTA integrity was linked to working memory.

Moreover, consistent with their dense interconnection and a largely shared biosynthesis, dopaminergic and noradrenergic brain regions’ integrity were positively related, and correlated with medial temporal lobe volumes. Longitudinally, we found that older age was associated with more-negative change in SN– VTA and LC integrity (time point 1–time point 2; mean delay ∼1.9 years). Importantly, changes in LC integrity reliably predicted future episodic memory performance (at time point 3).

These findings support the feasibility of in-vivo indices for catecholaminergic integrity with potential clinical utility, given the degeneration of both neuromodulatory systems in several age-associated diseases. Moreover, they highlight differential roles of dopaminergic and noradrenergic neuromodulatory nuclei in late-life cognitive decline.

## Introduction

Our memory fades as we age [1]. On average, old age is characterized by impaired abilities to retain and manipulate information over shorter time scales—termed working memory [2–4]—and to recollect past experiences within their temporal and spatial context—called episodic memory [1,5,6]. On a neural level, senescent declines in memory have been linked to dopaminergic neuromodulation [7–9] and, more recently, also to noradrenergic neuromodulation [10–12]. The degeneration of catecholaminergic (i.e. dopaminergic and noradrenergic) systems also is a core feature of age-related pathologies, such as Alzheimer’s and Parkinson’s disease [11,13–18] that are characterized by amnestic impairments [19,20]. However, investigations disentangling the contribution of the two neuromodulators to human memory in aging and disease are scarce.

Neuromodulators are a group of neurochemicals synthesized in circumscribed subcortical nuclei from which they are released throughout the brain via widely branching axonal projections [21]. Dopaminergic neurons are based mainly in the midbrain substantia nigra–ventral tegmental area (SN–VTA) [22], whereas noradrenergic neurons are primarily found in the brainstem locus coeruleus (LC) [23].

There are several mechanistic accounts linking dopaminergic and noradrenergic neuromodulation to aging memory. Computational models posit that catecholamines modulate the input–output relation of neurons (i.e., gain change) which increases the signal-to-noise ratio in neural processing [24] and influences cognition [25–27]. Age-related neurodegeneration of dopaminergic and noradrenergic nuclei thus results in noisier neural information processing (i.e., gain reduction) [8]. Specifically, declining catecholaminergic drive with increasing age is hypothesized to lead to less distinctive cortical representations and senescent memory decline [8,28].

A second mechanism linking dopaminergic and noradrenergic neuromodulation to aging memory is their modulation of prefrontal processing [29]. Lateral prefrontal circuits can represent external stimuli in the absence of sensory stimulation, even in the face of distractors, by means of persistently firing delay cells [30]. Catecholaminergic inputs orchestrate recurrent activity in delay cell circuits that is essential for higher-order cognitive functions, such as working memory [31]. Specifically, the stimulation of dopaminergic D1-receptors and noradrenergic α_2a_-receptors boosts prefrontal delay activity with an inverted-u dose–response curve [12]. Age-related memory deficits, in turn, have been associated with reduced delay cell firing, which could be partially restored by catecholaminergic drugs [32–35].

Finally, dopamine and noradrenaline modulate hippocampal long-term potentiation (LTP) and long-term depression (LTD) [36–40], which are critical for synaptic plasticity and memory. Initial accounts proposed a ventral tegmental area–hippocampal circuit by which neuromodulatory inputs facilitate the consolidation of salient experiences [36,39,41]. Interestingly, more recent investigations indicate that while the SN–VTA and LC both project to the dorsal hippocampus, the latter sends denser inputs [42–44]. LC neurons also produce dopamine as biosynthetic precursor of noradrenaline [45], and can co-release both catecholamines to modulate hippocampal synaptic plasticity and memory [42,43,46–48]. Older age is characterized by impaired hippocampal plasticity [49,50], likely exacerbated by deficient catecholaminergic innervation from the LC and SN–VTA [51].

Taken together, dopaminergic and noradrenergic neuromodulation mechanistically sculpts senescent memory via several pathways, including gain modulation [8,10], frontal delay activity[30,32], and hippocampal synaptic plasticity [36,47]. Notably, these mechanisms are specified for dopaminergic *and* noradrenergic neuromodulation (gain modulation [24]; delay activity [12]; synaptic plasticity [47]). However, research sampling a broad array of cognitive tasks to identify *unique* associations with late-life memory is lacking. That is, while animal research has demonstrated considerable overlap of catecholamines at the neural level, the question remains how much dopaminergic and noradrenergic nuclei overlap in their association with behavior.

Comparative studies of catecholaminergic systems in humans were long hampered by technical challenges in reliable non-invasive assessments of the small subcortical nuclei [52,53]. However, recent advances in high-resolution magnetic resonance imaging (MRI) reveal the SN–VTA [17,54] and LC [55,56] as focal hyperintensity on MRI scans. Multimodal post-mortem validation studies suggest this hyperintensity as a marker for catecholaminergic neurons [57–59]. Neuromelanin, a catecholamine-derived paramagnetic pigment accumulating in the LC and SN–VTA [60,61], presumably contributes to the MRI contrast of the nuclei [56,58,59,62,63]. However, also other factors likely play a role, such as the large cellular bodies of catecholaminergic neurons [64,65] that results in a high abundance of ions and water protons [65] as well as a lower macromolecular fraction [66,67]. Importantly, first in-vivo studies suggest an association between the MRI intensity of catecholaminergic nuclei and their functionality [58]. Furthermore, investigations in clinical populations confirm the validity of the imaging approach [17,68– 72].

Catecholaminergic nuclei are among the first brain structures to accumulate pathologies in age-associative diseases like Parkinson’s and Alzheimer’s [13,73,74] and show severe degeneration over the course of these diseases [11,14,16]. In line with this, LC [68–70] and SN–VTA [17,71,72] imaging using dedicated MRI sequences (i.e., Magnetization Transfer [MT], Fast Spin Echo [FSE]) reveals reduced hyperintensities in patients relative to controls. In healthy lifespan samples, initial cross-sectional evidence reveals a negative quadratic relationship between age and catecholaminergic hyperintensity [75–77], whereby lower contrast with advancing age might be linked to impending pathology [68,78,79]. Taken together, recent advances in imaging techniques open the door for comparative non-invasive assessments of catecholaminergic nuclei integrity, which are sensitive for age and disease-related changes [17,54,56,71,72].

In the present study, we took advantage of these new imaging approaches to attempt to disentangle the relative contribution of declining dopaminergic and noradrenergic neuromodulation to aging cognition. We repeatedly assessed cognitive performance and high-resolution MRI in large samples of younger and older adults across multiple time points [80–83]. Furthermore, we leveraged latent-variable modeling of multimodal imaging [84,85] and comprehensive cognitive assessments [85,86] to decrease measurement error and increase generalizability [87]. In sum, the goal of this study was to extend our knowledge about the respective roles of dopaminergic and noradrenergic neuromodulation in late-life memory decline.

## Results

### Locus coeruleus and substantia nigra–ventral tegmental area intensity shows high agreement across imaging modalities

We applied a validated semi-automatic procedure [88–92] to extract intensity information of catecholaminergic nuclei from different imaging sequences (FSE, MT+, and MT–; see Methods and Figure S2)—by standardizing MRI intensity in the LC and SN–VTA based on the intensity in an adjacent white matter reference region [55,56,58] (see Figures 1 and 2). Based on earlier post-mortem validations [57–59,69], we interpret individual differences in standardized MRI intensity as proxy for the integrity of catecholaminergic nuclei. Next, we leveraged an established factor structure [88] to integrate intensity information across hemispheres for each imaging sequence and age group (see Figure S3).

**Figure 1.**
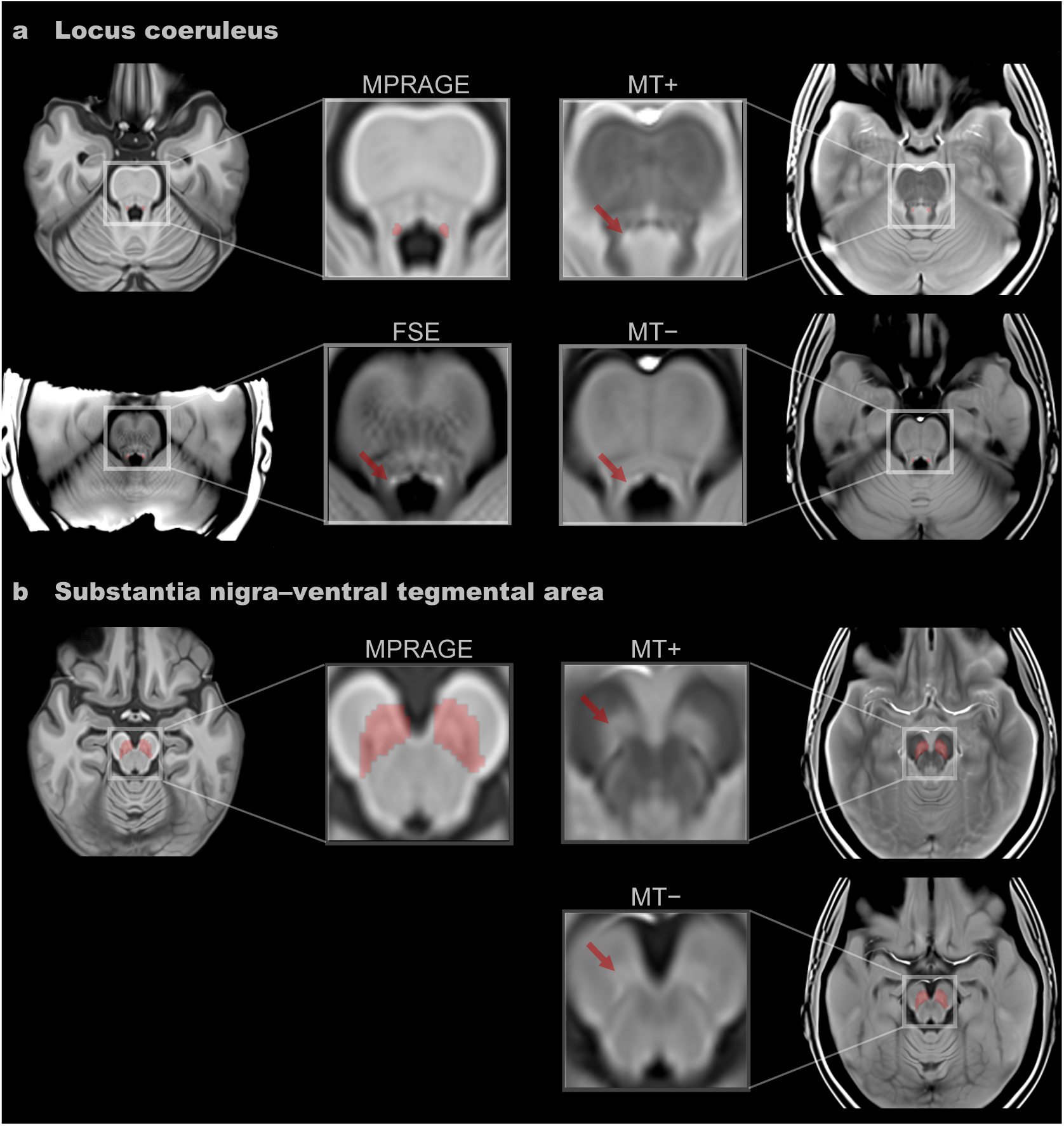
LC and SN–VTA-sensitive magnetic resonance imaging sequences. Hyperintensities corresponding to the (**a**) LC and (**b**) SN–VTA are indicated by red arrows on axial slices of group templates based on a Fast Spin Echo (FSE) sequence, and a Magnetization Transfer sequence, acquired once with a dedicated magnetic saturation pulse (MT+) and once without, resulting in a proton density image (MT–). A Magnetization Prepared Gradient-Echo (MPRAGE) sequence template shows the location of a previously-established LC [68] and SN–VTA [58] volume of interest (red overlays). Note that our FSE sequence only covered the brainstem. All templates were registered to MNI152 0.5mm linear space and are available at [94]. Templates were estimated across age groups. LC, locus coeruleus; SN–VTA, substantia nigra–ventral tegmental area.

**Figure 2.**
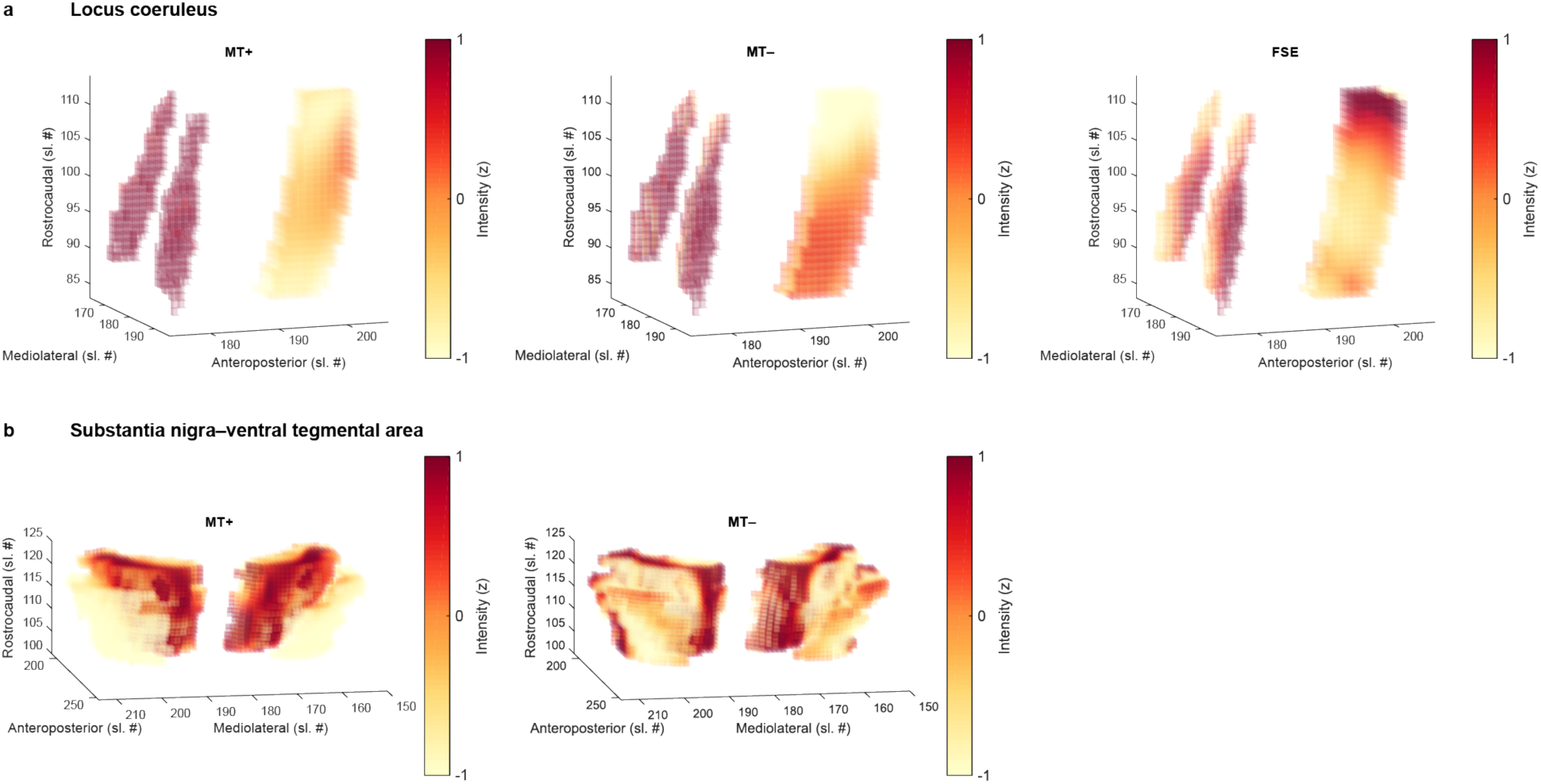
Normalized intensity in the LC, SN–VTA, and corresponding white-matter reference regions. Hyperintensities corresponding to the (**a**) LC and (**b**) SN–VTA are evident in group templates based on a Fast Spin Echo (FSE) sequence, and a Magnetization Transfer sequence, acquired once with a dedicated magnetic saturation pulse (MT+) and once without, resulting in a proton density image (MT–). LC and reference volumes of interest are taken from [68], SN–VTA and reference volumes of interest are based on [58]. The reference regions are located anterior of the LC and SN–VTA, respectively. In the visualization, they can be seen as rectangular shape to the right of the LC (i.e., the two red columns) as well as in front of the SN–VTA (i.e., the red curved shape). Note that our FSE sequence only covered the brainstem. Templates were estimated across age groups. LC, locus coeruleus; SN– VTA, substantia nigra–ventral tegmental area.

Previous in-vivo studies of catecholaminergic nuclei relied on different imaging approaches (mostly MT+ and FSE; [17,54–56]), but there are few comparisons between these MRI sequences, limiting cross-study comparability. Contrasting LC and SN–VTA estimates, we found strong differences across MRI sequences in their average intensity, Δ*χ*²(*df* = 2) = 693.55; *p* < 0.001 for older adults’ LC; Δ*χ*²(*df* = 1) = 657.37; *p* < 0.001 for older adults’ SN–VTA. That is, standardized to a reference region, catecholaminergic nuclei appeared brightest in the MT+, followed by the FSE, and finally the MT– sequences, for older adults’ LC (mean [standard error]): MT+, 25.816 (0.304); FSE, 20.144 (0.37); MT–, 6.425 (0.194); for older adults’ SN–VTA: MT+, 19.523 (0.235); MT–, 2.979 (0.135). Crucially, despite these mean differences, intensities were highly correlated across imaging modalities, *r* =0.43–0.621; *p* < 0.001 for older adults’ LC; *r* =0.503; *p* < 0.001 for older adults’ SN–VTA (see Figure 3 and S6–8), indicating that these sequences provide convergent information about the same underlying construct (i.e., catecholaminergic nuclei).

**Figure 3.**
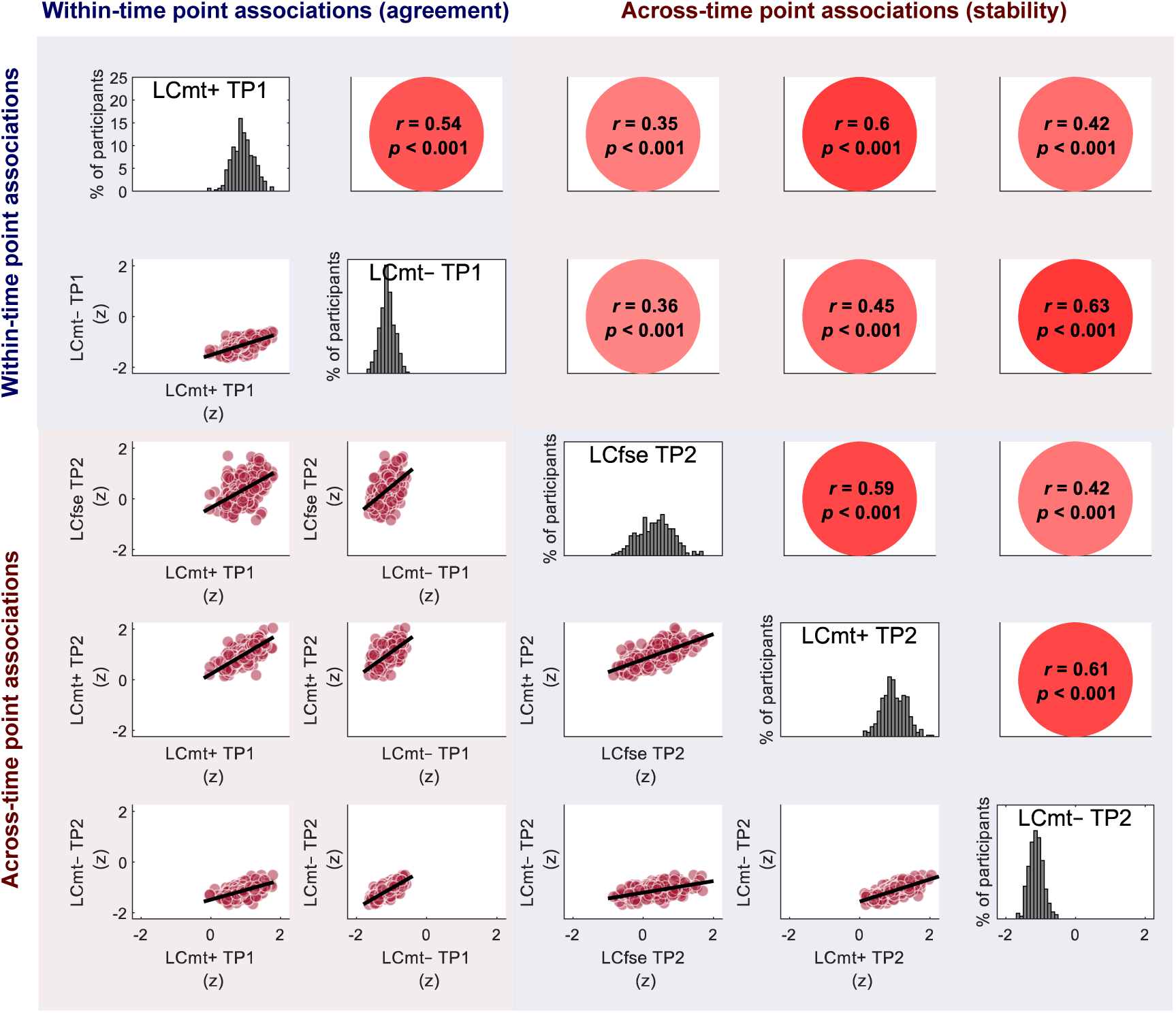
Younger and older adults’ LC intensities are correlated across imaging modalities—a marker for their agreement—and across time points—a marker for their stability. Visualized data are based on the statistical model depicted in Figure S14. For the same analyses using SN–VTA data, see Figures S18 and S20. Note, the diagonal shows LC intensity, standardized across all sequences and time points, to facilitate comparing intensity distributions. Imaging sequences included a Fast Spin Echo (FSE) sequence, and a Magnetization Transfer sequence, acquired once with a dedicated magnetic saturation pulse (MT+) and once without, yielding a proton density image (MT–). LC, locus coeruleus; TP, time point; SN–VTA, substantia nigra–ventral tegmental area.

**Figure 4.**
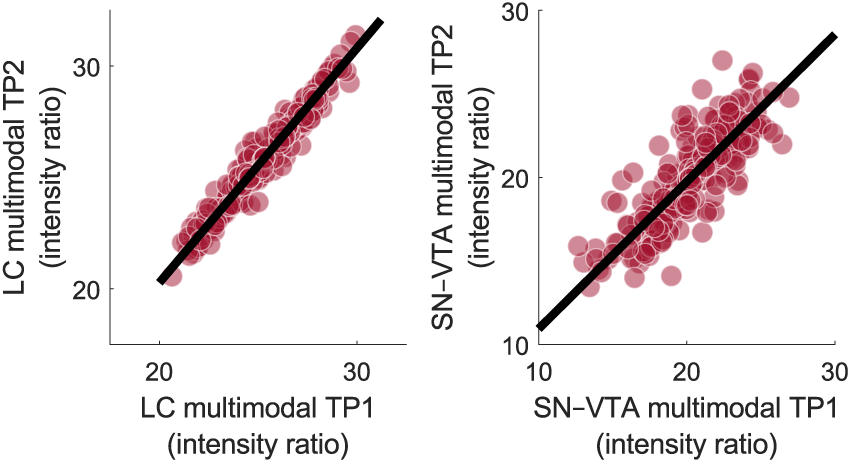
Multimodal LC and SN–VTA estimates show high stability over time points (mean delay ∼ 1.9 years), *r* = 0.88; *p* < 0.001 for younger and older adults’ LC (test against mean modality-specific stability, i.e., *r* = 0.615, *z* = 11.713; *p* < 0.001); *r* = 0.67; *p* < 0.001 for younger and older adults’ SN–VTA (test against mean modality-specific stability [i.e., *r* = 0.448], *z* = 5.837; *p* < 0.001). LC, locus coeruleus; SN–VTA, substantia nigra–ventral tegmental area.

We thus aggregated the information shared across imaging modalities by estimating multimodal latent factors expressing LC integrity and SN–VTA integrity (see Figure S4; for comparable approaches and a discussion, see [85] and [84,93]). Such latent variables capture the commonalities across scan modalities while removing the modality-specific measurement error and hence increase statistical power to detect true effects [87]. Model visualizations, model fit, and younger adults’ findings are reported in the supplementary information (see Figures S3–4, S6–7, and [94]).

Taken together, we extracted the intensity of the LC and SN–VTA from different MRI sequences sensitive for catecholaminergic nuclei. We found a high agreement in intensities across imaging modalities and thus aggregated across sequences to obtain individual integrity estimates for the two catecholaminergic nuclei.

### Multimodal locus coeruleus and substantia nigra–ventral tegmental area integrity factors show high stability over time

Making use of the longitudinal nature of this dataset, we next explored the stability of our integrity estimates over time (TP1–TP2; mean delay ∼1.9 (SD 0.7) years), as a proxy for their reliability [95]. Longitudinal studies investigating the reliability of imaging sequences for catecholaminergic nuclei are sparse. Thus, as a reference, we first assessed the stability of the modality-specific intensity factors and found evidence for an intermediate stability, for younger and older adults’ LC: MT+, *r* = 0.6; *p* < 0.001; MT–, *r* = 0.63; *p* < 0.001; for younger and older adults’ SN–VTA: MT+, *r* = 0.66; *p* < 0.001; MT–, *r* = 0.18; *p* = 0.17 (see Figures 3 and S14, S18, S20; FSE sequence only available for TP2 [see Methods]). For comparable analyses using intensity values extracted from the pontine reference (LC) and crus cerebri reference (SN–VTA) regions, see Figures S23 and S24.

If our multimodal integrity factors remove modality-specific measurement error [85], we should observe a higher stability across time points for the multimodal as compared to the modality-specific factors.

Importantly, our multimodal LC and SN–VTA factors indeed evinced a higher stability, pointing to the benefits of the multimodal imaging approach, *r* = 0.88; *p* < 0.001 for younger and older adults’ LC (test against mean modality-specific stability [i.e., *r* = 0.615], *z* = 11.713, *p* < 0.001); *r* = 0.67; *p* < 0.001 for younger and older adults’ SN–VTA (test against mean modality-specific stability [i.e., *r* = 0.448], *z* = 5.837; *p* < 0.001) [96].

### Locus coeruleus and substantia nigra–ventral tegmental area are associated with different aspects of late-life memory performance

Next, we cross-sectionally probed the behavioral implications of interindividual differences in LC and SN–VTA integrity, using data of TP2. For this, we leveraged a comprehensive cognitive battery and a previously established factor structure [86] to aggregate performance across several working memory, episodic memory, and fluid intelligence tasks and capture their shared variance on a latent level (see Figure S9). We observed strong age differences in each of the cognitive domains (see Figure 5), older relative to younger adults (mean age difference [standard error]): working memory, –2.265 (0.309); episodic memory, –2.287 (0.309); fluid intelligence, –2.073 (0.295); all *p* < 0.001. Importantly, however, when we added estimates for the catecholaminergic nuclei to the model, we found that higher LC and SN–VTA integrity were related to better performance (i.e., less age-related cognitive impairments) across domains, Δ*χ*²(*df* = 3) = 25.11; *p* < 0.001 for older adults’ LC; Δ*χ*²(*df* = 3) = 7.86; *p* = 0.049 for older adults’ SN–VTA (see Figure S10).

**Figure 5.**
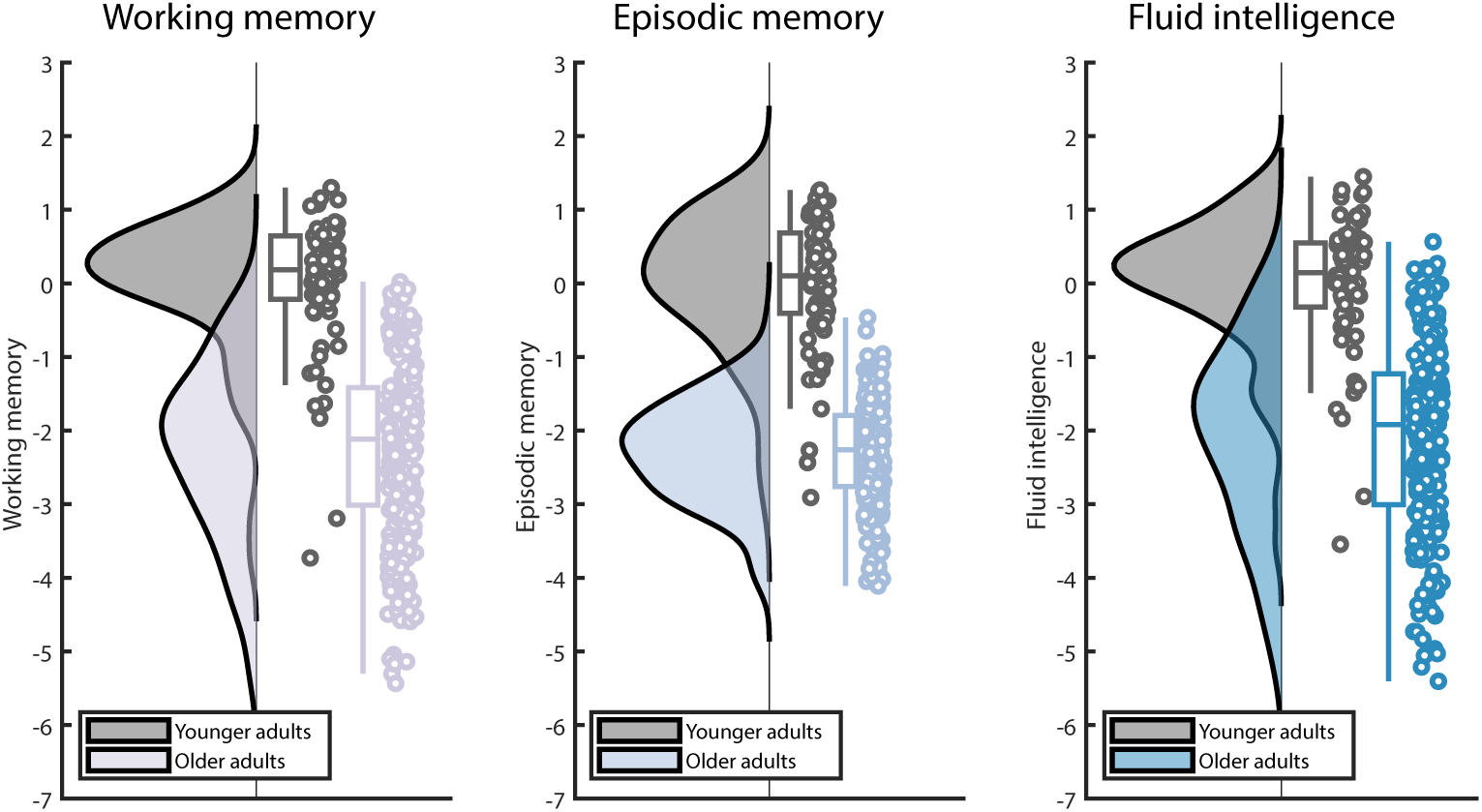
Cross-sectional age differences in working memory, episodic memory, and fluid intelligence factors at TP2 (for the statistical model, see Figure S9). Older adults show lower performance relative to younger adults across cognitive domains (mean age difference [standard error]): working memory, –2.265 (0.309); episodic memory, – 2.287 (0.309); fluid intelligence, –2.073 (0.295); all *p* < 0.001. Raincloud plots based on [100]. For visualizations of episodic memory and working memory performance over time points 1–3, see Figure S27–28.

Dopaminergic and noradrenergic neuromodulatory centers are densely interconnected [38,97]. In addition, dopamine is the biosynthetic precursor of noradrenaline [45], and indeed we detected a positive association between the structural metrics of the two neuromodulatory nuclei, *r* = 0.25; *Δ*χ²(*df* = 1) = 5.75; *p* = 0.017 for older adults (cf. Figure 6 and see S10). However, while the LC and SN–VTA were positively coupled, they, differed in their association with late-life cognition, *Δ*χ²(*df* = 3) = 15.66; *p* = 0.001 for older adults.

**Figure 6.**
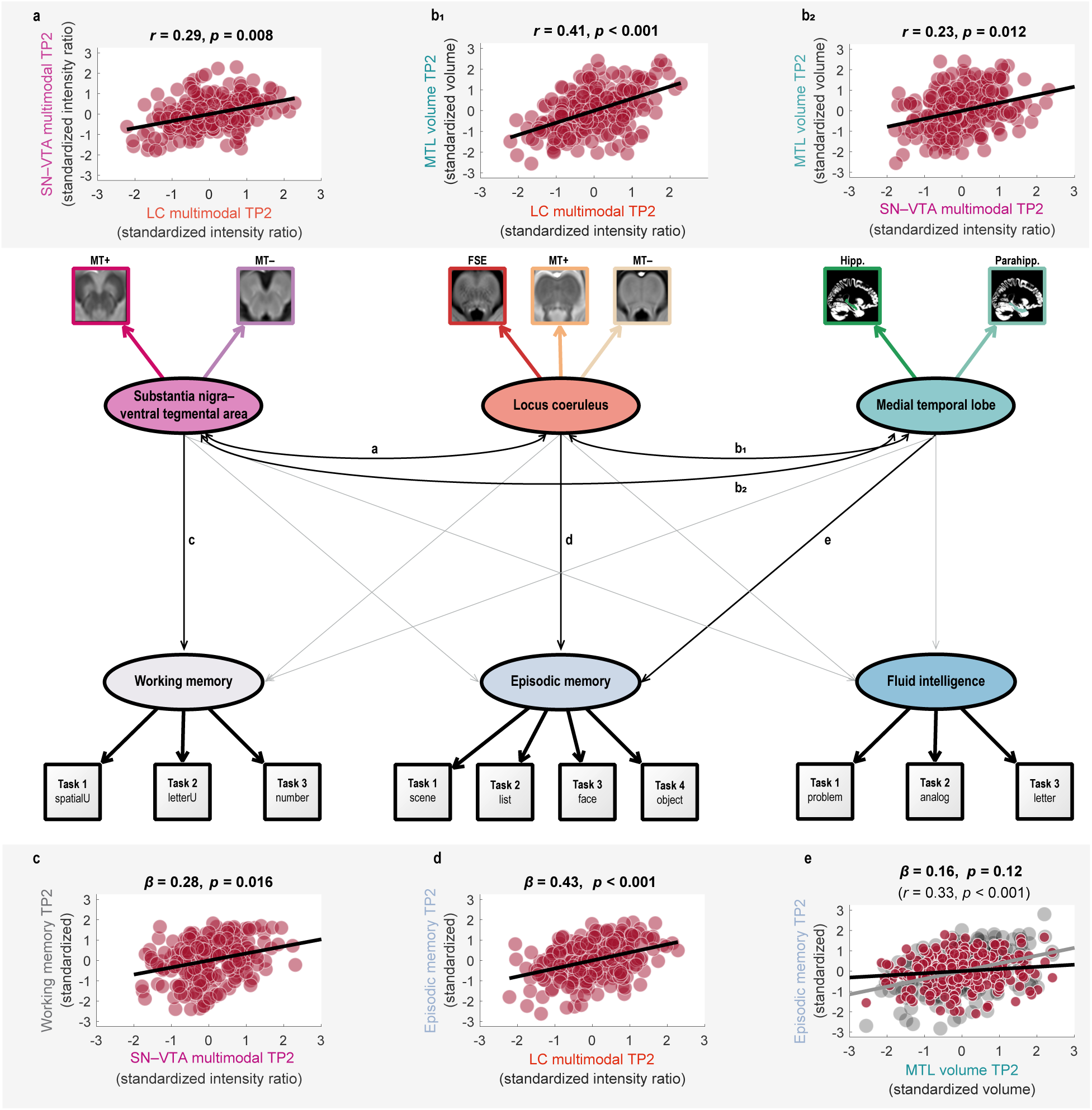
Schematic pictorial rendition of a structural equation model probing the interrelation of catecholaminergic nuclei and medial temporal lobe volume (paths **a** and **b**) and their unique associations with late-life cognition (paths **c–e**) in older adults at TP2. Note that covariances among cognitive factors, intercepts, and error terms are omitted for clarity. For the full statistical model, see Figure S13. Task*, cognitive paradigm (indicator; [square]) for the respective cognitive domain (latent factor; [ellipse]); One-headed arrows, regression; Double-headed arrow, correlation. Note that the brightness of paths indicates their significance. Medial temporal lobe volumes only demonstrate a reliable association with episodic memory in a correlational model (see grey line in scatter plot **e**; cf. Figure S12), but not when controlling for catecholaminergic neuromodulation (see black line in scatter plot **e**). Locus coeruleus (LC) and substantia nigra–ventral tegmental area (SN–VTA) factors are derived from a Fast Spin Echo (FSE) sequence, and a Magnetization Transfer sequence, acquired once with a dedicated magnetic saturation pulse (MT+) and once without, resulting in a proton density image (MT–). Hipp, hippocampus volume; Parahipp, parahippocampus volume.

Follow-up analyses showed that higher LC integrity was related to better episodic memory performance, *r* = 0.49; *Δ*χ²(*df* = 1) = 21.44; *p* < 0.001 for older adults (cf. Figure 6 and see S10) and that this association was specific: That is, LC’s relation to cognition differed across domains (episodic memory, working memory, and fluid intelligence), *Δ*χ²(*df* = 2) = 10.64; *p* = 0.005 for older adults. Moreover, the LC– episodic memory association was stronger than that of the SN–VTA and episodic memory, *Δ*χ²(*df* = 1) = 6.63; *p* = 0.01 for older adults. Taken together, our findings show a general (i.e., task and imaging sequence independent), yet specific relation of LC and late-life episodic memory performance, corroborating and extending earlier work [68,69,88,98,99].

In contrast to the LC, higher SN–VTA integrity correlated with better late-life working memory, *r* = 0.28; *Δ*χ²(*df* = 1) = 6.76; *p* = 0.009 for older adults (cf. Figure 6 and see S10). There was a numerical tendency for differential associations of SN–VTA integrity with performance in each of the cognitive domains (working memory, episodic memory, and fluid intelligence), *Δ*χ²(*df* = 2) = 5.73; *p* = 0.057 for older adults. However, we did not observe reliable differences in the relation of the two neuromodulatory nuclei to working memory, *Δ*χ²(*df* = 1) = 2.01; *p* = 0.156 for older adults. In sum, our findings suggest a differential association of the two neuromodulatory systems with late-life memory performance. While episodic memory was associated with LC integrity, SN–VTA integrity was related to working memory.

Note that our model (see Figure S10) specified correlations between the latent catecholaminergic and cognitive factors. That is, it probed associations of one neuromodulatory system with cognition irrespective of the other neuromodulatory system. Thus, we additionally specified a second, statistically equivalent model in which we searched for unique associations with cognition for each catecholaminergic system while controlling for the respective other system, using a multiple regression approach (see [85] for a comparable approach; see Figure 6 and S11). Crucially, we again detected reliable LC–episodic memory, *β* = 0.5; *Δ*χ²(*df* = 1) = 19.55; *p* < 0.001, and SN–VTA–working memory associations, *β* = 0.28; *Δ*χ²(*df* = 1) = 6.05; *p* = 0.014, for older adults, underlining differential relations to cognition of the two neuromodulatory nuclei.

Younger and older adults showed comparable associations between neuromodulatory integrity and memory performance (test for age group differences in (1) the LC–episodic memory association: (Δ*χ*²(df = 1) = 0.43; *p* = 0.512; and (2) the SN–VTA–working memory association: Δ*χ*²(df = 1) = 0.28; *p* = 0.594). Model visualizations, model fit, and younger adults’ findings are reported in the supplementary information (see Figures S10 and S11 and [94]).

### Locus coeruleus and substantia nigra–ventral tegmental area are associated with memory performance over and above medial temporal lobe volumes

Via direct projections, the LC and SN–VTA release catecholamines in memory-relevant brain regions, such as the medial temporal lobe [42,44,47], which improves retention performance [36,38,42,44,46,47]. Abnormally phosphorylated tau, an indicator of neurodegenerative diseases such as Alzheimer’s, starts to accumulate early in life in catecholaminergic nuclei [13,101–103]. With advancing age, abnormal tau also appears in projection targets of the neuromodulatory nuclei, like the medial temporal lobe [13,102,104,105], which may facilitate degeneration [68,69,105,106]. Thus, as a control analysis, we also incorporated medial temporal lobe volumes in our models linking catecholaminergic nuclei and cognition (see Figure S12 and S13).

In a correlational model, we observed that the integrity of both catecholaminergic nuclei was positively associated with medial temporal lobe volumes, *r* = 0.41; *Δ*χ²(*df* = 1) = 27.45; *p* < 0.001 for older adults’ LC; *r* = 0.23; *Δ*χ²(*df* = 1) = 6.29; *p* = 0.012 for older adults’ SN–VTA (cf. Figure 6 and see S12), potentially indicating neuroprotective catecholaminergic effects [107,108]. In addition, higher medial temporal lobe volumes were related to better late-life episodic memory performance, *r* = 0.33; *Δ*χ²(*df* = 1) = 14.22; *p* < 0.001 for older adults, in line with its role in memory processing [109].

Importantly, when we specified a second, multiple-regression model that searches for unique effects, we found that the LC was still reliably associated with episodic memory performance, *β* = 0.43; *Δ*χ²(*df* = 1) = 11.96; *p* < 0.001 for older adults, while medial temporal lobe volumes were not, *β* = 0.16; *Δ*χ²(*df* = 1) = 2.46; *p* = 0.117 for older adults (see Figure 6 and S13), when accounting for the respective other regions (e.g., controlling for SN–VTA and medial temporal lobe volume when evaluating the association between LC and episodic memory). Similarly, the SN–VTA was related to working memory after controlling for medial temporal lobe volumes and LC integrity, *β* = 0.28; *Δ*χ²(*df* = 1) = 5.8; *p* = 0.016 for older adults. For comparable analyses that are based on intensity values averaged across the LC or SN–VTA, see Table S 5.

Taken together, our results suggest a robust association of catecholaminergic nuclei and memory that cannot be fully accounted for by medial temporal lobe volume.

### Longitudinal changes in locus coeruleus integrity predict future episodic memory performance

Cross-sectional studies point to late-life differences in catecholaminergic nuclei [69,75–77,88,110], however, longitudinal data showing within-person changes are scarce. Thus, we combined imaging data of the two time points (TP1–TP2; mean delay ∼1.9 years) to test for individual changes in LC and SN– VTA integrity estimates in later life [87,111].

First, we observed that change in the catecholaminergic nuclei was correlated across imaging modalities (MT+, MT–; no longitudinal FSE data available), suggesting that the different MR sequences pick up a common underlying construct (i.e., change in catecholaminergic nuclei), *r* = 0.16; *Δ*χ²(*df* = 1) = 6.09; *p* = 0.014 for older adults’ LC; *r* = 0.13; *Δ*χ²(*df* = 1) = 5.91; *p* = 0.015 for older adults’ SN–VTA (see Figure S16 and S21). Thus, we again integrated across sequences to retrieve multimodal latent change factors for LC and SN–VTA integrity (see Figure S17 and S22). For both catecholaminergic systems, we found reliable individual differences in change, *Δ*χ²(*df* = 1) = 6.09; *p* = 0.014 for older adults’ LC; *Δ*χ²(*df* = 1) = 5.91; *p* = 0.015 for older adults’ SN–VTA. However, there was no reliable mean change at the group level in either nucleus, *p*s > 0.1 in older adults. That is, we observed that older adults differed from one another in how their LC and SN–VTA changed over time—while some older adults showed increases in intensity ratios, others showed decreases (see Figure 7; cf. [69,75]). Control analyses indicated, that changes in neuromodulatory integrity were not associated with the spatial positions from which intensity ratios were sampled at time point 1 and 2, making movement in the scanner an unlikely explanation for individual differences in change (see Figure S32).

**Figure 7.**
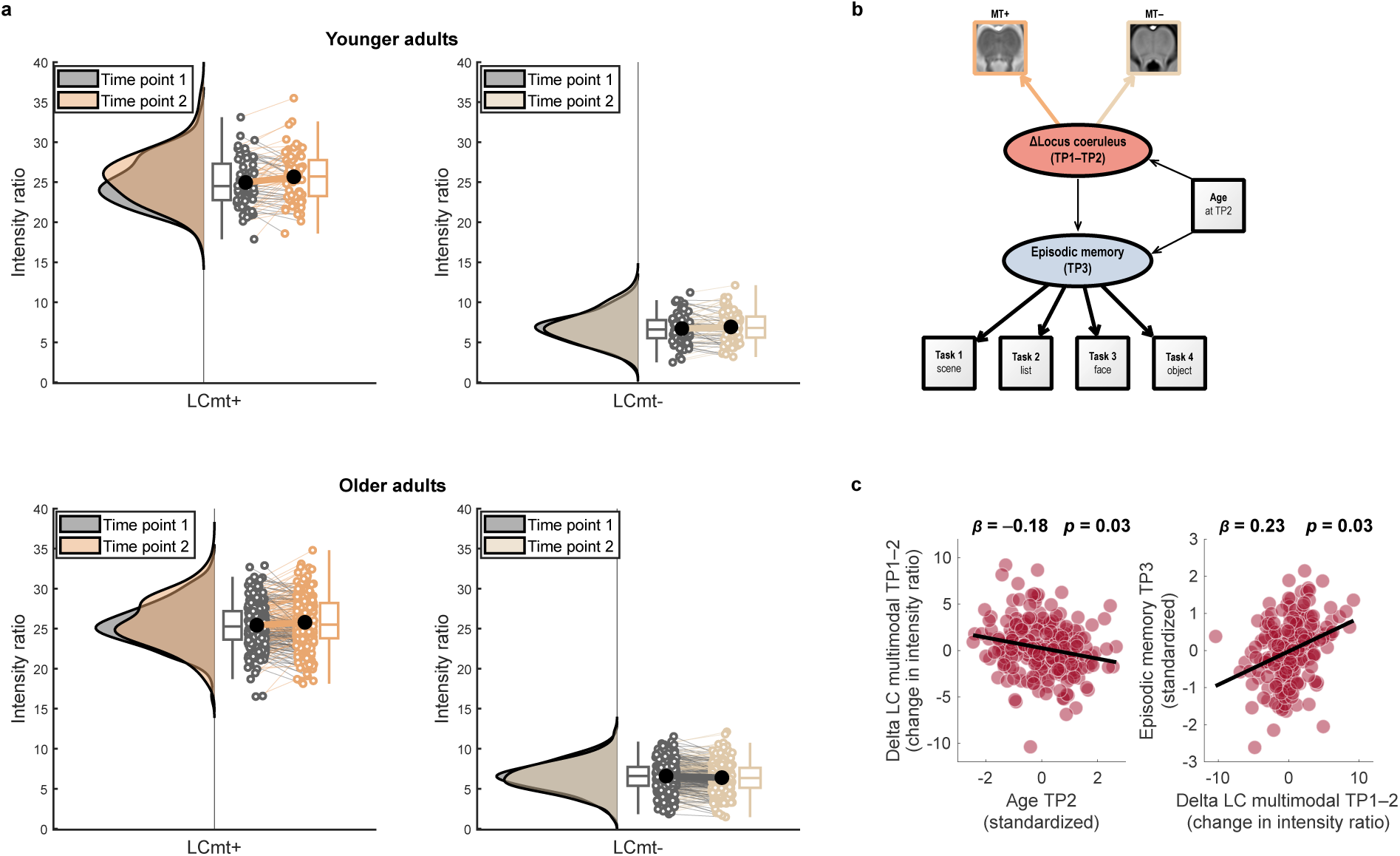
Longitudinal changes in LC intensity ratios and their association with age and future memory performance in older adults. **a**, Numerically, older adults show more negative average change in LC intensity across time points as compared to younger adults. MRI sequences include a Magnetization Transfer sequence, acquired once with a dedicated magnetic saturation pulse (MT+) and once without, resulting in a proton density image (MT–). For the Fast Spin Echo-sequence, only cross-sectional data are available. **b,** Schematic depiction of the structural equation model probing the association of longitudinal change in multimodal LC integrity with future episodic memory performance, accounting for chronological age. For the full model, see Figure S29. **c**, Scatter plots showing (1) more negative LC change in older adults of higher age and (2) a prediction of future memory performance based on LC change (controlling for chronological age). For comparable analyses using SN–VTA and working memory data, see Figures S30 and S31. Raincloud plots based on [100]. LC, locus coeruleus; SN–VTA, substantia nigra–ventral tegmental area.

Finally, to test the behavioral implications of these late-life changes in catecholaminergic nuclei, we probed whether our neural change model (TP1–TP2) could be used to predict future cognition (at TP3; using the previously established cognitive factor structure, cf. Figure S9). We additionally included chronological age (at TP2) as a predictor, to test and account for potential age effects in rate of decline (see Figure S29 and S30).

Note that we here related changes in catecholaminergic nuclei to subsequent cognitive performance (at TP 3), as our analyses indicated task-specific memory changes (see methods and supplementary results). In line with a late-life degeneration of neuromodulatory centers [7,10,11,69], we found more negative change in the LC and SN–VTA with increasing age, *β* = –0.18; *Δ*χ²(*df* = 1) = 4.81; *p* = 0.028 for older adults’ LC; *β* = –0.29; *Δ*χ²(*df* = 1) = 3.95; *p* = 0.047 for older adults’ SN–VTA. Moreover, we observed that changes in LC integrity predicted subsequent episodic memory performance over and above chronological age, *β* = 0.23; *Δ*χ²(*df* = 1) = 4.73; *p* = 0.03 for older adults’ LC (see Figure 7); *β* = 0.27; *Δ*χ²(*df* = 1) = 1.55; *p* = 0.213 for older adults’ SN–VTA (see Figure S31). Taken together, our results are in line with the proposition that a decline of the LC in later life is associated with diminished episodic memory performance [10,69,88,112].

## Discussion

This study sought to disentangle the effects of declining dopaminergic and noradrenergic neuromodulation on late-life memory. We took advantage of a multimodal imaging protocol and extensive cognitive assessments across several time points to contrast the behavioral implications of LC and SN–VTA integrity.

Our findings indicate that different imaging approaches for catecholaminergic nuclei (FSE, MT+, MT–) show a high agreement. Thus, we used latent-variable modeling to integrate across MRI modalities and retrieve multimodal LC and SN–VTA integrity factors that demonstrated a markedly increased reliability compared with their individual components. After establishing reliable in-vivo integrity proxies, we probed their associations with late-life cognition. We used an extensive neuropsychological test battery and a previously identified factor structure to demonstrate that these two catecholaminergic systems, although positively coupled, differ in their relationship three domains of aging cognition: episodic memory, working memory, and fluid intelligence.

We observed a general (i.e., task and imaging sequence independent), yet specific association of LC integrity and late-life episodic memory performance (i.e., stronger for episodic memory as compared to working memory and fluid intelligence). By contrast, SN–VTA integrity was linked to better working memory. Remarkably, both associations remained reliable even after accounting for the respective other neuromodulatory system and a key node in the memory network, the medial temporal lobe, suggesting robust effects. Corroborating this interpretation, associations between catecholaminergic integrity and late-life memory performance were qualitatively unchanged when including participants’ age, education, and sex as covariates (see *Supplementary results*).

Leveraging the longitudinal nature of this dataset, we also investigated late-life changes in the LC and SN–VTA over a delay of approximately two years. A principal finding was that older age was associated with more negative change in each catecholaminergic system, in line with the late-life degeneration of neuromodulatory centers. Moreover, we showed that changes in the LC predicted future episodic memory performance over and above chronological age and education (see *Supplementary results*). Taken together, this study suggests dopaminergic and noradrenergic neuromodulation play domain-specific roles in determining the trajectory of cognition in later life and provides novel insights into the neural basis of human senescent memory decline.

The loss of dopaminergic neuromodulation has long been recognized as crucial determinant of late-life cognitive deficits [7–9]. More recently, technical advances have also facilitated studies of the noradrenergic LC [56] that long seemed unattainable [52,53]. We used two types of imaging sequences (FSE, MT+) validated on postmortem specimen of the LC [57] and SN–VTA [58,59]. Neuromelanin, an insoluble catecholamine-derived pigment that traps metals and exhibits paramagnetic properties, is thought to contribute to the hyperintensity of catecholaminergic nuclei [17,56,58,59,62,63]. Moreover, the high density of water protons and paramagnetic ions in large catecholaminergic neurons has also been proposed as source of their MRI contrast [65,66]. In line with this, we provide the first quantification of the LC and SN–VTA based on a proton density-weighted sequence (i.e., without dedicated magnetization transfer preparation pulse; MT–). Magnetization transfer imaging studies frequently estimate a ratio score based on sequences with and without dedicated preparation pulse (i.e., MT+ and MT–) [113]. However, the sensitivity of our MT– sequence for the LC and SN–VTA suggests that this ratio would reduce the detectability of these nuclei. Notably, we observed that despite differences in mean contrast, LC and SN– VTA hyperintensities were correlated across all imaging modalities (FSE, MT+, MT–), suggesting that they provide convergent information about the same underlying construct (i.e., catecholaminergic nuclei).

We thus leveraged our multimodal approach to estimate latent factors for catecholaminergic nuclei integrity based on the commonalities across imaging sequences while removing modality-specific measurement error [84,85]. Moreover, using data from both imaging time points, we show that semiautomatic analyses of the intensity of catecholaminergic nuclei have high reliability, especially when multimodal assessments are available [114].

The retention of salient experiences is enhanced by catecholamine release in the hippocampus which facilitates synaptic plasticity and memory [36,38,47]. While the SN–VTA has long been attributed as source of the memory-enhancing dopaminergic inputs [36,39,44], recent findings point to a denser innervation by the LC that can provide noradrenergic as well as dopaminergic signals [42,46,47]. Here we compared the association of the two catecholaminergic centers with an extensive set of tasks that are thought to depend more (episodic memory) or less (working memory) on hippocampal processing [109,115]. We observed that LC integrity was specifically related to late-life episodic memory performance (as compared to working memory or fluid intelligence) and that this association was stronger than the SN–VTA–episodic memory relationship. While we observed comparable associations between neuromodulatory integrity and memory performance when comparing age groups, the lower sample size for younger adults [57] may have limited our ability to detect effects in this group alone. Mechanistically, our finding of an LC–episodic memory association might be explained by a catecholaminergic modulation of hippocampal synaptic plasticity [42,46,47], but our data cannot rule out other memory-related mechanisms, such as gain modulation [8,24–27] and prefrontal delay activity [12,30]. Our observations are supported by a series of large-scale in-vivo imaging studies that showed reliable LC– cognition associations in aging [69,88,98,99], and particularly with episodic memory [69,88,116]. In addition, they concord with a recent report linking anteromedial and superior substantia nigra intensity to attentional performance [117], a cognitive concept overlapping with working memory [25].

Animal research suggests also a noradrenergic role in attentional processes [12,25,118], particularly in tasks that require attentional re-orienting [119,120]. By contrast, the working memory indicator tasks used in the current study (e.g., number n-back) require participants to hold information active in mind and may thus depend less on noradrenergic neuromodulation (but see [32]). Interestingly, time-resolved measures associated with phasic locus coeruleus activity (pupil dilation, P-300 event-related potential [26,121–123]) show consistent associations with individual differences in attentional performance [124,125], calling for more multimodal research.

Cross-sectional studies point to late-life differences in catecholaminergic nuclei [75–77,88,110]. Here we provide the first characterization of late-life longitudinal changes in the LC and SN–VTA. In general agreement with extrapolations from cross-sectional data, we found more negative change in both catecholaminergic systems with increasing age. Subcortical neuromodulatory centers like the LC and the SN–VTA are among the first sites to accumulate pathology in age-associated diseases like Alzheimer’s [13] and Parkinson’s [73], and show severe degeneration with disease progression [16,126]. In combination with earlier work suggesting lower catecholaminergic contrast with advancing age might be linked to impending pathology [68,71,78,79,117], our findings may indicate subthreshold pathological processes in a subset of our older participants. This interpretation is supported by our prediction of poorer future memory performance in individuals with more negative LC change [112]. Accordingly, recent meta-analyses demonstrate the efficacy of noradrenergic treatments in improving cognitive symptoms in Alzheimer’s disease [127]. Mirroring its clinical potential, MRI-indexed catecholaminergic integrity has been suggested as useful tool for stratifying patients suffering for neurodegenerative diseases in clinical trials that include noradrenergic treatments [56,128]. A subset of older participants also showed LC and SN–VTA intensity increases over time, which might indicate higher intracellular proton density [66], potentially linked to the activity-related volume increase of catecholaminergic cells [11,15,129,130].

The present study highlights the utility of multimodal longitudinal assessments of catecholaminergic nuclei to elucidate the neurobiological basis of senescent memory decline. We dissociated the roles of the noradrenergic LC and dopaminergic SN–VTA in late life cognition. While the former showed robust associations with current and future episodic memory performance, the latter showed a relationship to better working memory performance. These findings point to differential roles of dopaminergic and noradrenergic neuromodulation in late-life cognition with potential implications for age-associated diseases like Alzheimer’s and Parkinson’s [13,69,73]. Accurate longitudinal assessments of catecholaminergic nuclei may constitute promising early markers for predicting cognitive decline in later life.

## Methods

### Study design and participants

Data were collected as part of the Berlin Aging Study-II (BASE-II), an ongoing longitudinal study that investigates neural, cognitive, physical, and social conditions related to successful aging (for more information, see https://www.base2.mpg.de/en, references [80–83], and the supplementary methods). Cognitive performance was assessed in three time periods (TP1–TP3) between 2013 and 2020 (TP1: 2013–2015; TP2: 2015–2016; TP3: 2018–2020) with a mean duration between cognitive assessments of 2.246 years (TP1–2; SD: 0.603) and 2.917 years (TP2–TP3; SD: 0.438).

A subset of BASE-II participants also underwent magnetic resonance imaging (MRI). Eligible participants had no history of neurological or psychiatric disorders, or head injuries, and did not take medication that may affect memory function. Imaging data were collected in two time periods (TP1, TP2) in temporal proximity to the cognitive assessments (mean delay between MRI waves 1.894 years; SD: 0.656). Participants were only considered for further analyses if at least one type of imaging sequence sensitive for dopaminergic or noradrenergic neuromodulatory centers was available (see below). For TP1, this corresponds to 288 participants out of a total of 488 participants with imaging data, whereas for TP2 this corresponds to 320 out of 323 participants with imaging data. Thus, our analyses included a total of 320 individual participants. While not all imaging sequences were available for all participants (see Table 1), 203 participants have relevant MRI data for both TP1 and TP2.

**Table 1:** Overview of participants with available imaging data

| Magnetic resonance imaging sequence | Time period | Number of participants with relevant imaging data | Total number of participants with imaging data |
| --- | --- | --- | --- |
| Magnetization Transfer (MT+) *and* Proton Density (MT–) | 1 | 288 | 448 |
| Magnetization Transfer (MT+) *and* Proton Density (MT–) | 2 | 260 | 323 |
| Fast Spin Echo (FSE) | 1 | 0 | 448 |
| Fast Spin Echo (FSE) | 2 | 316 | 323 |
| Fast Spin Echo (FSE) *and* Magnetization Transfer (MT+) *and* Proton Density (MT–) | 1 | 0 | 448 |
| Fast Spin Echo (FSE) *or* Magnetization Transfer (MT+) *and* Proton Density (MT–) | 1 | 288 | 448 |
| Fast Spin Echo (FSE) *and* Magnetization Transfer (MT+) *and* Proton Density (MT–) | 2 | 256 | 323 |
| Fast Spin Echo (FSE) *or* Magnetization Transfer (MT+) *and* Proton Density (MT–) | 2 | 320 | 323 |

*Note:* Relevant imaging data refers to participants with a whole-brain T<sub>1</sub>-weighted (MPRAGE) sequence as well as a sequence sensitive for dopaminergic or noradrenergic neuromodulatory centers (Magnetization Transfer (MT+), Proton Density (MT–) or Fast Spin Echo (FSE)). By contrast, the total number of participants with imaging data reflects all participants that underwent MRI, irrespective of sequence type. The Magnetization Transfer and Proton Density sequences were acquired in succession in each scan session (i.e., identical sequence, acquired once with and one without dedicated MT preparation pulse). Thus, they are grouped in the table.

The final sample (n = 320) included 69 younger adults (22 female; mean age (SD): 32.705 (3.884) years [at TP2]) and 251 older adults (91 female; mean age (SD): 72.414 (4.045) years [at TP2]). Sample descriptives are reported in Table 2.

**Table 2:** Summary of sample descriptives for younger and older adults

|  | Younger adults<br>(n = 69; 22 female) |  | Older adults<br>(n = 251; 91 female) |  |
| --- | --- | --- | --- | --- |
|  | Mean (SD) | Range | Mean (SD) | Range |
| Age<br>(years; at time period 2) | 32.705 (3.884) | 25.414–40.203 | 72.414 (4.045) | 62.534–83.162 |
| Education (years)<br>(available for n = 285) | 14.242 (2.519) | 10–18 | 14.08 (2.924) | 7–18 |
| Mini Mental State<br>Examination<br>(available for n = 204) | – | – | 28.613 (1.336) | 24–30 |

*Note:* Mini Mental State data was assessed around TP3, during a medical examination at a different visit as part of the longitudinal Berlin Aging Study-II. *SD*, Standard Deviation.

The cognitive and imaging assessments were approved by the institutional review boards of the Max Planck Institute for Human Development and the German Psychological Society (DGPS), respectively. Participants provided written informed consent and were reimbursed for their participation.

### Cognitive data assessment

At TP1–TP3, cognition was tested using a comprehensive computerized battery probing key cognitive functions. Performance was assessed in small groups of four to six participants. Cognitive test sessions lasted approximately 3.7 h and included 16 tasks (at TP2), of these, three measured working memory, four episodic memory, and three fluid intelligence [131]. While the exact composition of the cognitive assessments changed across waves, the same tasks were used to measure working memory, episodic memory and fluid intelligence at TP1–TP3. Only older adults were tested at TP3. For a detailed task description, please refer to [131–133], below we provide a brief overview of the measures relevant to the current analyses.

### Working memory assessment

#### *Spatial Updating* (abbreviated as spatial-*u*)

Participants were shown a display of two to three 3 × 3 grids, in each of which a blue dot was presented in one of the nine tiles. Participants were asked to memorize the locations of the blue dots and mentally update them according to shifting operations that were indicated by arrows appearing below the dots. Six updating operations were required before the 3 × 3 grids reappeared and participants indicated the end position of the blue dots (by mentally combining their start position and the six shifting operations). We used the number of correct placements as an indicator of working memory performance [131,133].

#### *Letter Updating* (abbreviated as letter-*u*)

Participants were shown a sequence of 7–13 letters. Once the presentation ended, they were asked to report, in correct order, the last three letters that were shown. The number of correctly reported letters was used as a measure of working memory performance [131].

#### *Number N-back* (abbreviated as *number*)

Three digits (1–9) were presented consecutively in three adjacent cells, followed by the next sequence of three digits. Participants indicated by button press whether the currently presented digit matched the digit shown three steps before [131,133]. We took participants’ accuracy as indicator of their working memory performance.

### Episodic memory assessment

#### *Scene encoding* (abbreviated as *scene*)

Participants incidentally encoded 88 scene images by performing indoor/outdoor judgments on each image. The encoding phase was followed by an old/new recognition memory test, which included confidence judgments. Recognition memory was tested after a delay of approximately 2.5 hours and served as episodic memory performance index (hits – false alarms) [85,88,131].

#### *Verbal learning and memory task* (abbreviated as *list*)

Participants first learned a 15-word list that was presented via headphones. The task was composed of five learning trials each followed by a free recall period in which participants entered the words they remembered via keyboard (trial 1–5; recall of learning list). After these initial learning–recall cycles, participants were presented an interference list and their delayed recall and recognition memory was assessed. The sum of correctly recalled words during the learning–recall cycles (trial 1–5) served as episodic memory measure [85,88,131].

#### *Face–profession task* (abbreviated as *face*)

Participants studied 45 pairs of face images and profession words. The tasks consisted of an incidental encoding phase, a 2-minute distraction phase, and finally a recognition memory task including old, new as well as rearranged face–profession pairs. Corrected recognition memory scores for rearranged pairs was used as the performance index.

#### *Object–location task* (abbreviated as *object*)

Participants encoded the location of 12 digital photographs of real life objects on a 6 × 6 grid. After encoding, the objects reappeared next to the grid and participants were instructed to reproduce their correct location by placing the items in the grid. The sum of correct placements served as index of episodic memory.

### Fluid intelligence assessment

#### *Practical problems* (abbreviated as *problem*)

Participants were sequentially presented eighteen items depicting everyday problems (e.g., the hours of a bus timetable, a subway map etc.), in order of ascending difficulty. For each of these problems, five response alternatives were provided and participants selected the correct option by clicking on it. We took the sum of correctly solved problems as measure of fluid intelligence [131,134].

#### *Figural analogies* (abbreviated as *analog*)

Participants were instructed to draw analogies. They were presented with twenty-two items in ascending difficulty that followed the format *A* is to *B* as *C* is to *X*? Below each item, five response alternatives were presented and participants selected the correct option by clicking on it. The sum of correctly given answers served as index of fluid intelligence [131,134].

#### *Letter series* (abbreviated as *letter*)

Participants were shown twenty-two series of five letters, each ending with a question mark (e.g., c-e-g-i-k-?). Each series followed a simple rule (e.g., +1, –1, +2; or +2, –1), with increasing difficulty. Below each letter series, five response alternatives were presented and participants selected the correct option by clicking on it. The sum of correct responses served as fluid intelligence measure [131,134].

All tasks included practice blocks to familiarize participants with the instructions. Note that these tasks have previously been used to estimate latent factors of working memory, episodic memory, and fluid intelligence [131] (also see [85,88,135]).

### Magnetic resonance imaging data assessment

To investigate the associations of dopaminergic and noradrenergic integrity with late-life cognition, younger and older participants underwent 3T-MRI at TP1 and TP2 (MAGNETOM TIM Trio, Siemens Healthcare). Only those sequences used in the current analyses are described below. The imaging protocol included three scans sensitive for the SN–VTA and LC—a Fast Spin Echo sequence (FSE; sometimes also called Turbo Spin Echo [TSE]), and a Magnetization Transfer sequence, acquired once with a dedicated magnetic saturation pulse (MT+) and once without, resulting in a proton density image (MT–). Moreover, a Magnetization Prepared Gradient-Echo (MPRAGE) sequence, comparable to the ADNI protocol (www.adni-info.org), was collected to facilitate coregistration to standard space and to estimate volumes for regions of interest. Moreover, the MPRAGE sequence was used during acquisition to align the FSE sequence perpendicularly to the plane of a participant’s brainstem. Note that for some participants specific absorption rate (SAR) limits were exceeded during the FSE acquisition, as reported previously [88] (also see [55,136]). Sequence parameters are reported in Table 3.

**Table 3:** Overview of magnetic resonance imaging sequences

| Magnetic resonance imaging sequence | Acquisition matrix; Slices (orientation; distance factor) | Voxel size (x, y, z; mm) | Repetition time (TR; ms) | Echo time (TE; ms) | Flip angle (°) |
| --- | --- | --- | --- | --- | --- |
| Magnetization Transfer (MT+)<sup>1</sup> | 192 × 256 × 48 (axial; –) | 1 × 1 × 3 | 38 | 5.5 | 10 |
| Proton Density (MT–)<sup>2</sup> | 192 × 256 × 48 (axial; –) | 1 × 1 × 3 | 38 | 5.5 | 10 |
| Fast Spin Echo (FSE)<sup>3</sup> | 440 × 512 10 (axial; 20 %) | 0.5 × 0.5 × 2.5 | 600 | 11 | 120 |
| Magnetization Prepared Gradient-Echo (MPRAGE)<sup>4</sup> | 256 × 256 × 192 (sagittal; –) | 1 × 1 × 1 | 2500 | 4.77 | 7 |

*Note:* All sequences were acquired with a standard 32-channel head coil.
<sup>1</sup>: Magnetization Transfer Contrast (MTC) option enabled (MT pulse: 1200 Hz off-resonance, 16 ms [110]); 3-D sequence
<sup>2</sup>: Magnetization Transfer Contrast (MTC) option disabled; 3-D sequence
<sup>3</sup>: Each FSE acquisition included four online averages and yielded two brainstem images. The values extracted from these images were aggregated offline. SAR limits resulted in a slight variation in the number of slices [88]; 2-D sequence; sometimes also called Turbo Spin Echo (TSE).
<sup>4</sup>: Inversion time (TI; ms): 1100; 3-D sequence

### Magnetic resonance imaging data analysis

We applied a previously established semi-automatic analysis procedure to extract individual LC and SN– VTA intensity values from structural imaging data (for a detailed description and validation, see [88], for applications, see [89,90,137]; for an independent validation see [92]). The following procedure was performed separately for TP1 and TP2 imaging data.

### Template construction and standardization

First (step 1), scans of each scan modality (MPRAGE, FSE, MT+ and MT–) were iteratively aligned across participants using a template-based procedure implemented in Advanced Normalization Tools [ANTs] (v. 2.3 [138,139]; *antsMultivariateTemplateConstruction,* 6 iterations, including *N4BiasFieldCorrection*). A schematic visualization of the procedure is included in the supplementary methods. Before template construction, MPRAGE and MT– scans were resampled to 0.5 mm isometric resolution (ANTs’ *ResampleImage*). Moreover, to facilitate template construction, participants’ native space FSE scans were aligned to their template-space MPRAGE scans (*antsRegistrationSyNQuick*) [88]. Native space MT+ scans were aligned to resampled MT– scans to account for potential movement effects between scan acquisitions (*antsRegistrationSyNQuick*). After their alignment, MT– and MT+ scans were submitted to a common multimodal template construction, while FSE and MPRAGE scans each were used to generate a brainstem and whole-brain template, respectively.

Next (step 2), modality-specific group templates (MPRAGE, FSE, MT+ and MT–) were linearly and non-linearly coregistered (*antsRegistration*) to standard space (MNI-ICBM 152 linear, 0.5 mm [140]).

Specifically, templates with a sensitivity for catecholaminergic nuclei (FSE, MT+, MT–) were first standardized to the whole-brain MPRAGE template (using a coregistration mask). Next, the MPRAGE template was coregistered to MNI space and the transformations were applied to the other templates (FSE, MT+, MT–; *antsApplyTransforms*). To improve coregistration accuracy, whole-brain templates (MPRGAE, MT–, MNI) were skull stripped before alignment using the FMRIB Software Library [141]. Finally (step 3), all transformation matrices were concatenated and applied to individual participants’ scans to bring them from native to MNI space in a single step (*antsApplyTransforms*).

### Semi-automatic intensity assessment

To extract the intensity values of catecholaminergic nuclei, in standard space, individual scans were masked with binary volumes of interest using Statistical Parametric Mapping toolbox version 12 (SPM12) [142] in Matlab (MathWorks). In particular, for the LC, we applied a previously established high-confidence consensus mask [68]. For the SN–VTA, we relied on a previously established mask that was based on manual tracings in template space [58]. Inter-participant comparisons of arbitrality scaled MRI intensity values require that intensity values are normalized within participants [56]. Thus, we also masked scans (FSE, MT+, MT–) with volumes of interest in potine [68] and midbrain [58] reference areas, in line with earlier research [55,58,88]. Note that the fourth ventricle, which is in close proximity to the LC, appears hyperintense in our MT+ scans. Thus, to rule out that the hyperintensity of the ventricle could interfer with automatized LC assessments, we generated a sample-specific ventricle volume-of-no-interest (based on the MT– group template), which we removed from MT+ and MT– scans before value extraction (templates and ventricle mask are available from [94]). Within the masked scans (FSE, MT+, MT–), we then automatically searched for the voxel of highest intensity in the LC, SN–VTA, and reference regions. Next, for each participant, spatially-resolved intensity ratios for the LC and SN–VTA were computed per hemisphere (left, right) on a slice-by-slice basis using the following formula [55,88]:

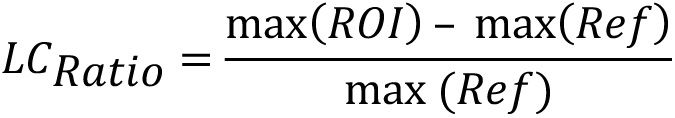

Where max(*ROI*) denotes the peak intensity for a given slice in the LC or SN–VTA regions of interest and max(*Ref*) indicates the peak intensity in the respective reference region. For the FSE modality, two scans were available per participant (see *Magnetic resonance imaging data assessment*) and we averaged the extracted intensity ratios within participants to obtain more stable estimates. For further analyses, for all modalities (FSE, MT+, MT–) the peak intensity ratio across the regions of interest (LC, SN–VTA) was calculated as an overall integrity measure [53,68,88]. Outlier values exceeding ± 3 standard deviations were dropped, while all other values were linearly scaled (× 100) to facilitate subsequent model estimation. Note that all analyses including LC or SN–VTA data were based on peak intensity ratios. That is, the peak intensity of catecholaminergic nuclei standardized using nearby white matter regions (not their raw intensity values). To facilitate readability, we will nonetheless use the term *intensity* in our description.

At acquisition, the FSE sequence was centered on the pons and contained fewer slices compared to the MT+ and MT– sequences. As evident in Figure 2 (a), the most rostral slices of our volume of interest (LC meta mask and reference mask) reach the edges of the brainstem template and include high intensity artifacts, which, however, are reliably excluded from analyses using the semiautomatic procedure (cf. peak detection above).

### Structural equation modeling

We used structural equation modeling (SEM) to evaluate inter- and intra-individual differences in catecholaminergic nuclei and their association with cognition using the Ωnyx software environment [143] and the lavaan R package [144]. All models used full information maximum likelihood (FIML) estimation to account for missing values. The adequacy of the reported models was evaluated using *χ*^2^-tests (i.e., an absolute fit index), as well as two frequently reported approximate fit indices: the root mean square error of approximation (RMSEA) and the comparative fit index (CFI). RMSEA values close to or below 0.06 and CFI values close to 0.95 or greater indicate good model fit [145,146]. Unless otherwise noted, multi-group models were fit, comprising submodels for younger and older adults. For this, invariance across age groups was evaluated by a hierarchical series of likelihood-ratio tests, probing group differences in (1) factor loadings (weak invariance), (2) indicator intercepts (strong invariance), and (3) residual variances (strict invariance) [147]. In case of longitudinal models, the same criteria were applied to test invariance across time [111]. After establishing adequate model fit and invariance, the significance of parameters of interest was evaluated using likelihood-ratio tests. That is, we created two nested models—in one, the parameter of interest was freely estimated, whereas in the other model it was fixed to zero. If fixing the parameter of interest to zero resulted in a drop in model fit, as evaluated using a likelihood-ratio test comparing the two nested models, this indicated the significance of the parameter [148]. We used an alpha level of .05 for all statistical tests. Statistical results with *p*-values between 0.05 and 0.1 are described as statistical trend. All analyses are based on two-sided statistical tests and did not include corrections beyond those mentioned in the respective sections. We assumed normally distributed cognitive and neural data, based on visualizations of data distributions (cf. e.g., Figures 3 and 5). In the following, cross-sectional models refer to models that include TP2 data only, while longitudinal models evaluate the change in parameters of interest over time (TP1–TP2 or TP1–TP3). Model code, visualizations, and output are available from [94].

### Cross-sectional cognitive models

We made use of a previously established factor structure [131] to integrate performance across several working memory, episodic memory, and fluid intelligence tasks (see *Cognitive data assessment*) and capture their shared variance on a latent level. Latent variables account for measurement error in the observed scores (cognitive tasks) and thus increase statistical power to detect true effects [87].

We added covariances between the latent working memory, episodic memory, and fluid intelligence factors, as performance across these cognitive domains had been shown to be correlated [131] (see Figure S9).

### Cross-sectional neural models

We also adapted a previously established factor structure [88] to capture LC and SN–VTA intensity on a latent level. Specifically, for each region (LC, SN–VTA) and scan modality (FSE, MT+, MT–), we used the left and right hemispheric peak intensity as observed scores to estimate a modality-specific integrity factor on a latent level. Note that our FSE sequence only covered the brainstem. Thus, we cannot obtain SN–VTA estimates for this scan modality. To test the agreement in integrity estimates across modalities (FSE, MT+, MT–), we added covariances among the modality-specific latent factors for each brain region (see Figure S3).

Using the model describe above, we found a high correspondence in the integrity estimates for each nucleus across modalities (see *Results*). Thus, in a second model, we introduced a multimodal integrity factor for the LC and SN–VTA that captures the commonalities across scan modalities while removing the modality-specific measurement error (for similar approaches and a detailed discussion, see [84,85]; see Figure S4). Finally, as dopamine and noradrenaline are products of the same biosynthesis pathway, with dopamine a direct precursor to noradrenaline [45], we evaluated the association of the multimodal LC and SN–VTA integrity factors by estimating their covariance.

### Cross-sectional neuro–cognitive models

After separately establishing models for our cognitive and neural measures that showed acceptable fit and invariance across age groups, we unified these models to probe associations between catecholaminergic nuclei and cognition [88]. In the unified neuro-cognitive model, we estimated covariances between the multimodal LC and SN–VTA integrity factors and working memory, episodic memory, and fluid intelligence (correlation model; see Figures S10 and S12). In addition, we specified a second neuro-cognitive model, in which regression paths were drawn from the latent neural to the cognitive factors (regression model; see Figures S11 and S13). The correlation model evaluates associations between the LC and cognition irrespective of those between the SN–VTA and cognition. By contrast, the regression model tests whether one region explains variance in the cognitive factors over and above the other region, thus providing complementary information [85].

### Longitudinal cognitive models

Making use of the repeated assessments of cognitive performance, we tested for late-life changes in working memory and episodic memory tasks over time (TP1–TP3). Note that cognitive data at TP3 was only available for older adults (see *Cognitive data assessment*). Thus, here we relied on single-group models that excluded younger adults. In particular, we specified a latent change score model [87,149] for each cognitive task. These latent change score models yield a latent slope factor for each task that expresses participants’ performance difference between TP1 and TP3 (see Figures S25 and S26).

Next, we tested whether there is a common latent factor of working memory or episodic memory change. If changes in performance were shared across tasks (i.e., task independent) of one cognitive domain, they could be captured on a higher order latent level (cf. the latent working memory and episodic memory factors in the cross-sectional cognitive model; Figure S9). To evaluate potential associations in the changes across working memory and episodic memory tasks, we added covariances among the task-specific slope terms (see Figures S25 and S26).

Most covariances across task-specific slope terms did not reach significance (see *Supplementary results*). Thus, we did not further attempt to capture task-independent changes in working memory or episodic memory on a latent level.

### Longitudinal neural models

#### Assessment of across-time stability

Our cross-sectional analyses of different MRI sequences sensitive for catecholaminergic nuclei (FSE, MT+, MT–) demonstrated a high agreement in integrity estimates across imaging modalities (see *Cross-sectional neural models* and *Results* Figure 3). Leveraging the longitudinal nature of this study, we additionally explored the stability of the modality-specific integrity estimates over time (TP1, TP2), as a proxy for their reliability [95]. Specifically, we started with a modality-specific SN–VTA and LC models for TP2 (see *Cross-sectional neural models* and Figure S3) and then appended the same variables for TP1. We introduced covariances linking modality-specific factors of TP1 and TP2 to evaluate the stability in integrity estimates over time (see Figures S14 and S18). Moreover, we allowed for correlated residuals over time, as suggested for longitudinal models [111].

Using the model describe above, we found a high stability of the modality-specific integrity estimates for each region across time (see *Results* Figure 3 and S20). Comparable to our cross-sectional analyses, we thus again added multimodal integrity factors for the LC and SN–VTA for each time point (TP1, TP2). If the multimodal integrity factors indeed remove measurement error [85], we should observe a higher stability across time points of the multimodal as compared to the modality-specific integrity factors. To test this hypothesis, we computed the covariance between the multimodal factors of TP1 and TP2 (see Figures S15 and S19). In addition, we directly compared the stability estimates, our reliability proxy, for the modality-specific and multimodal SN–VTA and LC factors. Note that the modality-specific and multimodal integrity models including TP1 and TP2 were fit across younger and older adults to obtain a single reliability proxy for the complete sample.

### Assessment of within-person changes

Previous cross-sectional research observed between-person age differences in catecholaminergic nuclei integrity [75,76], however, longitudinal studies that evaluate within-person changes in the SN–VTA and LC are scarce. Making use of the imaging data of both time points (TP1, TP2), we thus estimated changes in catecholaminergic nuclei using latent change score models [87,149] for each imaging modality (MT+, MT–). Note that the FSE sequence was only acquired at TP2; precluding change analyses (see Table 1). To reduce model complexity, we averaged intensity ratios across hemispheres for these models.

Comparable to our longitudinal cognitive analyses, we first evaluated whether the change in each region was shared across imaging modalities (i.e., sequence independent) by computing the covariance of the modality-specific slope terms (see Figures S16 and S21).

For the SN–VTA as well as the LC we observed that the changes were indeed correlated across imaging modalities (MT+, MT–). Thus, in a second set of models, we introduced a higher order multimodal slope factor to capture the shared variance across modality-specific slope factors (see Figures S17 and S22).

### Longitudinal neuro–cognitive models

In older adults, we found that SN–VTA and LC integrity changed across time (TP1–TP2; see *Longitudinal neural models*). To evaluate the behavioral implications of these changes, we leveraged a previously established cognitive factor structure [131,135] (see *Cross-sectional cognitive models*).

Specifically, we were interested in testing whether changes in catecholaminergic nuclei (TP1–TP2) could be used to predict future cognition (at TP3). For this, we unified our neural change models (see Figures S17 and S22) with models of working memory and episodic memory (at TP3; cf. Figure S9). In the unified model, we specified regression paths from the neural change factors to the cognitive factors (SN– VTA to working memory; LC to episodic memory). Finally, we added chronological age (at TP2) as an additional predictor to test (1) whether catecholaminergic nuclei explains future cognition over and above age and (2) how changes in catecholaminergic nuclei differ in old age (see Figures S29 and S30). Note that cognition at TP3 was only assessed for older adults. Thus, we specified single-group models excluding younger adults.

### Control analyses

Catecholaminergic neuromodulation influences neural processing in the medial temporal lobe, a key node of the memory network [36,38,42,44,46,47]. Moreover, catecholaminergic nuclei have direct projections to the medial temporal lobe [42,44,150,151] and their integrity has been linked to tau pathology in these areas [68–70]. Thus, as a control analysis, we additionally evaluated medial temporal lobe volumes (at TP2). This allowed us to compare our measures of catecholaminergic nuclei and their association with memory performance to those of a well-established player in the memory network [109].

### Voxel-based morphometry assessment

Whole-brain MPRAGE images were processed using the voxel-based morphometry pipeline in SPM12 in MATLAB [142,152]. First, images were segmented into distinct tissue classes (e.g., grey matter, white matter, cerebrospinal fluid) using SPM12’s Unified Segmentation procedure [153]. Next, a study-specific DARTEL template was created and segmented images were aligned to the template, followed by spatial normalization, modulation, and smoothing with a 2 mm full-width at half-maximum isotropic Gaussian kernel [154,155]. The resulting normalized, modulated, and smoothed grey matter images were used to derive region-of-interest volumes. Region-of-interest masks from the AAL3 atlas were applied to processed grey matter images using native SPM functions and the get_totals script (http://www0.cs.ucl.ac.uk/staff/g.ridgway/vbm/get_totals.m) to calculate volumes for the parahippocampal and hippocampal regions [156]. Outlier values exceeding ± 3 standard deviations were dropped, while all other values were linearly scaled (× 10,000) to facilitate subsequent model estimation. Before analyses, volume data were adjusted by dividing regional estimates by total intracranial volume.

### Cross-sectional neuro–cognitive models including voxel-based morphometry factor

Mimicking our integrity factors of catecholaminergic nuclei (see Figure S3), we aggregated across left and right hemispheric volumes to estimate latent parahippocampal and hippocampal factors for older adults. We specified a covariance between the two regions to evaluate their association (see Figure S5). We observed that parahippocampal and hippocampal factors were highly correlated (see *Supplementary results*). Thus, we introduced a higher-order medial temporal lobe factor to capture their shared variance, which we then included in our neuro-cognitive models (see *Cross-sectional neuro–cognitive models*).

Specifically, for older adults we estimated one correlation model and one regression model [85]. In each model, we compared the associations of catecholaminergic nuclei and cognition with the medial temporal lobe–cognition association (see Figures 6 and S12, S13). In addition, we quantified the interrelations of SN–VTA and LC integrity and medial temporal lobe volume by estimating their covariance.

### Data availability

The data that our results are based on are available from the BASE-II steering committee upon approved research proposal (see https://www.base2.mpg.de/en). For inquiries, please contact Dr. Ludmilla Müller, BASE-II project coordinator.

To facilitate comparability of study results, we share the group templates with sensitivity for catecholaminergic nuclei (FSE, MT+, MT–) in MNI 0.5 mm linear space (https://osf.io/eph9a/) [94]. The LC consensus volume of interest (LC meta mask and pontine reference mask) is available via https://osf.io/sf2ky/ [90].

We provide two synthetic datasets of simulated cases (n = 250) that follow the population described in our models, with the parameter values displayed in the model visualizations (generated using [143]). In combination with the model code (see below), these data allow reproduction of key results.

### Code availability

All statistical models that our inferences are based on and their outputs are available via: https://osf.io/eph9a [94]

Vector-graphics visualizations for all models are also available via OSF.

Additional custom code used for processing behavioral and neural data are available from the corresponding authors upon request.

## Acknowledgements

This article uses data from the Berlin Aging Study II (BASE-II), which was supported by the German Federal Ministry of Education and Research (Bundesministerium für Bildung und Forschung, BMBF) under grant numbers #16SV5536 K, #16SV5537, #16SV5538, and #16SV5837, #01UW070 and #01UW0808. Additional contributions (e.g., financial, equipment, logistics, personnel) are made from each of the other participating sites, i.e., the Max Planck Institute for Human Development (MPIB), Max Planck Institute for Molecular Genetics (MPIMG), Charite-Universiätsmedizin, University Medicine, German Institute for Economic Research (DIW), Humboldt-Universität zu Berlin, all located in Berlin, Germany, and University of Lübeck, and University of Tübingen, Germany. For further information about the BASE-II project, see https://www.base2.mpg.de/en

MW-B received support from the German Research Foundation (DFG, WE 4269/5-1) and the Jacobs Foundation (Early Career Research Fellowship 2017–2019).

MM’s work was supported by an Alexander von Humboldt fellowship and by National Institutes of Health grant R01AG025340.

SB was supported by a National Institutes of Health grant T32AG000037 and a National Science Foundation Grant DGE-1842487.

SD received funding from the National Science Foundation (DGE1418060).

The funders had no role in study design, data collection and analysis, decision to publish, or preparation of the manuscript.

We thank Dr. Clifford Cassidy for providing the substantia nigra and crus cerebri masks [58]. We are grateful to Dr. Ylva Köhncke and Dr. Sarah Polk for preprocessing the cognitive data.

We thank Sumedha Attanti for visually inspecting the VBM-processed segmentations.

