## Supplementary information for "The integrity of dopaminergic and noradrenergic brain regions is associated with different aspects of late-life memory performance"

**Title:**

*** Corresponding author:**

**Supplementary information:**

**Supplementary methods** ... 3

Sampling in the Berlin Aging Study-II ... 3

**Figure S1.** Flowchart depicting the participant selection for each time point and age group. ... 4

**Figure S2.** Overview of MRI template generation ... 5

**Supplementary results** ... 6

**Table S1.** Overview of model paths evaluated using likelihood ratio tests ... 6

Cross-sectional neural models: ... 10

**Table S2.** Model fit and invariance for cross-sectional neural models ... 10

**Figure S3**. Model 1.1.1 ... 11

**Figure S4**. Model 1.1.2 ... 12

**Figure S5**. Model 1.1.3 ... 13

**Figure S6**. Cross-sectional age differences in modality-specific LC factors. ... 14

**Figure S7.** Cross-sectional age differences in modality-specific SN–VTA factors ... 15

**Figure S8.** LC and SN–VTA intensities are correlated across imaging modalities (in older adults). 16

Cross-sectional cognitive models: ... 17

**Table S3**. Model fit and invariance for cross-sectional cognitive models ... 17

**Figure S9.** Model 1.2.1 ... 18

Cross-sectional neuro–cognitive models: ... 19

**Table S4**. Model fit and invariance for cross-sectional neuro–cognitive models ... 19

**Figure S10. Model 1.3.1. (a)** ... 20

**Figure S11. Model 1.3.1. (b)** ... 21

**Figure S12. Model 1.3.2. (a)** ... 22

**Figure S13. Model 1.3.2. (b)** ... 23

Cross-sectional neuro–cognitive model with mean intensity ratios ... 24

**Table S5**. Comparison of parameter estimates of cross-sectional neuro-cognitive models fit with peak and mean intensity ratio data ... 24

Longitudinal neural models: ... 25

**Table S5**. Model fit and invariance for longitudinal neural models ... 25

**Figure S14.** Model 2.1.1 ... 26

**Figure S15.** Model 2.1.2 ... 27

**Figure S16.** Model 2.1.3 ... 28

**Figure S17.** Model 2.1.4 ... 29

### Supplementary methods

#### Sampling in the Berlin Aging Study-II

The Berlin Aging Study-II (BASE-II) is a study of healthy aging—participants were cognitively unimpaired at baseline. Younger (20–35 years of age) and older participants (60–80 years of age) were enrolled at time point 1 and no new participants were entered afterwards. The study design and sampling are described in several recent publications [1–4]. The following description of the sampling procedure represents a verbatim quote from [2]:

*Only residents of the greater metropolitan area of Berlin, Germany, were eligible for participation in BASE-II. Potential participants were drawn from a pool of individuals originally recruited at the Max-Planck-Institute for Human Development as part of a number of earlier projects with a focus on neurocognition. Briefly, participant recruitment for these and other studies was based on advertisements in local newspapers and the public commuter transport system. This led to approximately 10 000 responders of whom 2875 were invited for an additional screening (either in-house or by telephone), leading to 2262 individuals eligible for inclusion in BASE-II, i.e. 79% of those who were initially invited. From these, we selected 2200 individuals to represent the BASE-II baseline cohort based on their age and sex as follows. A total of 1600 participants were assigned to an older subgroup aged between 60 and 80 years, whereas the remaining 600 individuals were assigned to a younger subgroup (serving as a reference population) aged between 20 and 35 years. By design, each age subgroup contains equal numbers of males and females. Some ageing-related changes, such as decline in perceptual speed, begin in early adulthood. At the same time, recent longitudinal studies indicate that average performance on other cognitive abilities, such as episodic memory, is relatively stable until about 60 years of age, and starts declining thereafter. Hence, we decided to start observing older adults at an age where most would show subsequent decline on most variables of interest. Comparisons with representative survey data from Berlin and Germany, ascertained via the SOEP questionnaire (see below), reveal that BASE-II participants are characterized by higher education and better self-reported health status than the general population of Berlin and Germany. In addition, BASE-II participants in the older subgroup report a significantly higher divorce/separation rate than participants in the age-matched reference populations. For convenience samples such as BASE-II this is a commonly observed phenomenon.*

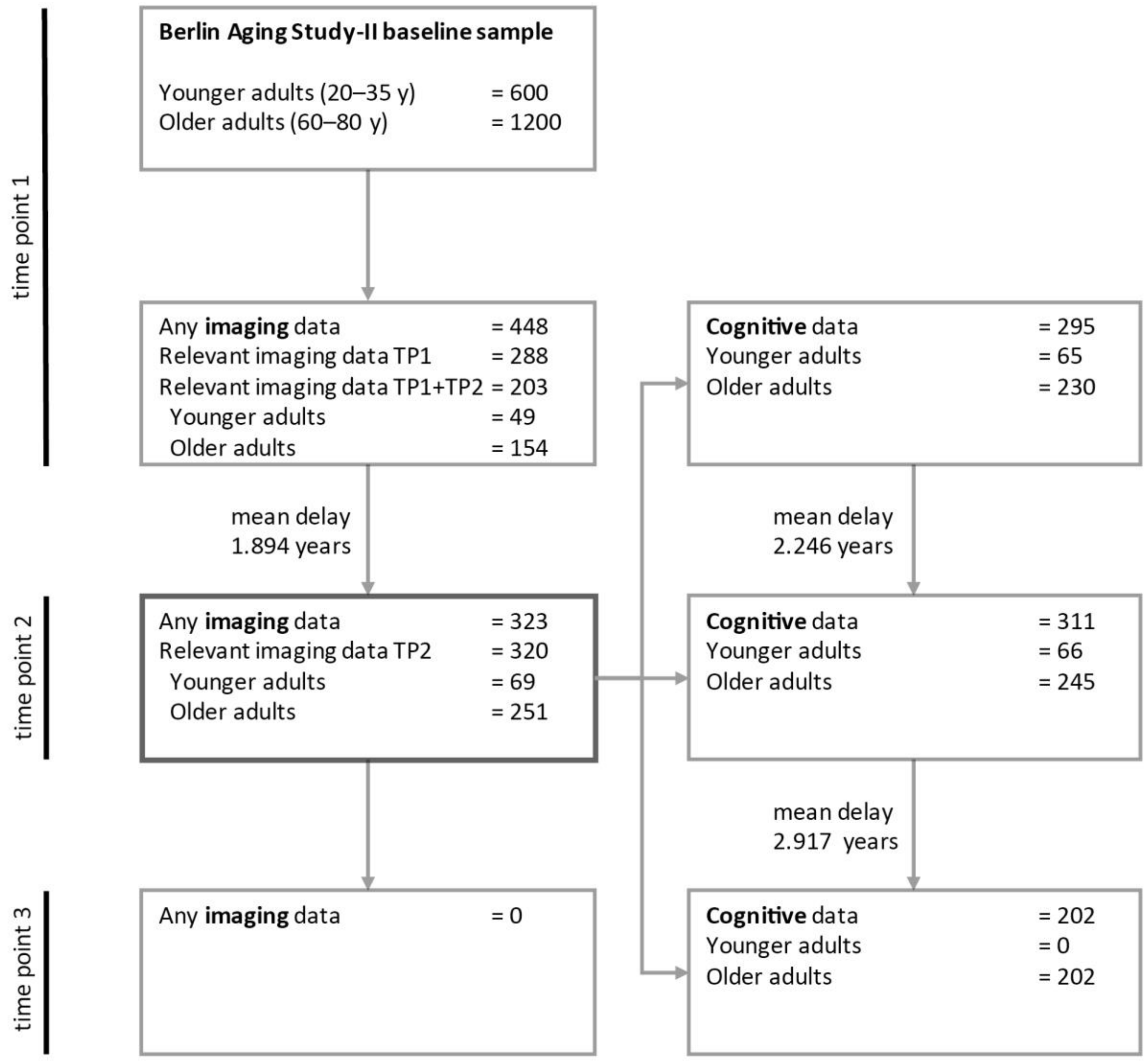

**Figure S1.** Flowchart depicting the participant selection for each time point and age group.

For general information on the eligibility criteria and recruitment procedure, see [2]. Note that analyses started with cross-sectional data (within time point 2; dark grey box) and were followed up by longitudinal analyses (brain changes from time point 1 to 2). Thus, all statistical models were restricted to the n = 320 participants with relevant imaging data at time point 2.

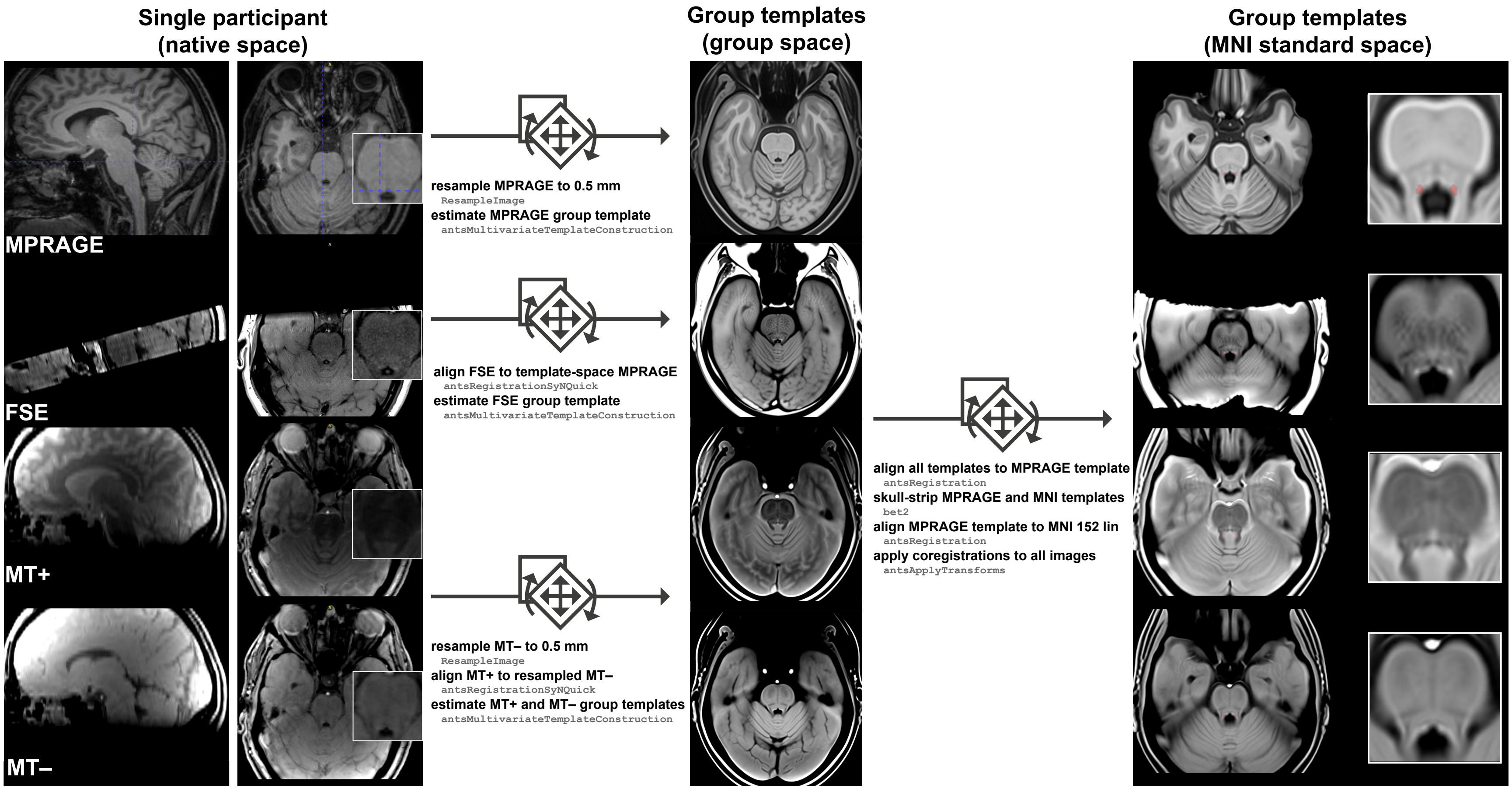

**Figure S2.** Overview of MRI template generation.

From left to right, images are transformed from native space via group template space to standard MNI 152 linear 0.5 mm space. The arrows indicate the transformations across spaces. Grey font indicates the corresponding functions. LC-related hyperintensities are evident across modalities (except MPRAGE) in single participant and group images. The blue crosshair marks the approximate location of the right locus coeruleus in native space. In standard space, the red overlay indicates the locus coeruleus volume of interest [5]. FSE, Fast Spin Echo; MT+, Magnetization Transfer; MT–, Proton Density.

### Supplementary results

**Table S1.** Overview of model paths evaluated using likelihood ratio tests

Result section:
**Locus coeruleus and substantia nigra–ventral tegmental area intensity shows high agreement across imaging modalities**

| Model | Tested group(s) (sample size) | Tested path(s) | Results | Interpretation |
|---|---|---|---|---|
| 1.1.1. | Multi-group model with YA, OA ($n_{YA}$ = 69; $n_{OA}$ = 251) | μLCfse'<br>μLCmt'<br>μLCnomt' | $\Delta\chi^2$(df = 2) = 693.55; $p$ < 0.001 | Mean locus coeruleus intensity differs across MRI sequences |
| 1.1.1. | Multi-group model with YA, OA ($n_{YA}$ = 69; $n_{OA}$ = 251) | μSNmt'<br>μSNnomt' | $\Delta\chi^2$(df = 1) = 657.37; $p$ < 0.001 | Mean substantia nigra–ventral tegmental area intensity differs across MRI sequences |
| 1.1.1. | Multi-group model with YA, OA ($n_{YA}$ = 69; $n_{OA}$ = 251) | γLCfse_LCmt'<br>γLCfse_LCnomt'<br>γLCmt_LCnomt' | $r$ = 0.61; $\Delta\chi^2$(df = 1) = 43.95; $p$ < 0.001;<br>$r$ = 0.43; $\Delta\chi^2$(df = 1) = 23.53; $p$ < 0.001;<br>$r$ = 0.62; $\Delta\chi^2$(df = 1) = 52.71; $p$ < 0.001 | Locus coeruleus intensity is correlated across MRI sequences (agreement) |
| 1.1.1. | Multi-group model with YA, OA ($n_{YA}$ = 69; $n_{OA}$ = 251) | γSNmt_SNnomt' | $r$ =0.503; $\Delta\chi^2$(df = 1) = 31.67; $p$ < 0.001 | Substantia nigra–ventral tegmental area intensity is correlated across MRI sequences (agreement) |

Result section:
**Multimodal locus coeruleus and substantia nigra–ventral tegmental area integrity factors show high stability over time**

| Model | | Tested path(s) | Results | Interpretation |
|---|---|---|---|---|
| 2.1.1. | Single-group model with YA_OA ($n_{YA–OA}$ = 320) | γLCmt_TP1_LCmt_TP2;<br>γLCnomt_TP1_LCnomt_TP2 | $r$ = 0.6; $\Delta\chi^2$(df = 1) = 40.32; $p$ < 0.001;<br>$r$ = 0.63; $\Delta\chi^2$(df = 1) = 52.57; $p$ < 0.001 | Locus coeruleus intensity is correlated within MRI sequences across time (stability) |
| 2.1.5 | Single-group model with YA_OA ($n_{YA–OA}$ = 320) | γSNmt_TP1_SNmt_TP2;<br>γSNnomt_TP1_SNnomt_TP2 | $r$ = 0.66; $\Delta\chi^2$(df = 1) = 45.84 $p$ < 0.001;<br>$r$ = 0.18; $\Delta\chi^2$(df = 1) = 1.88; $p$ = 0.17; | Substantia nigra–ventral tegmental area intensity is correlated within MRI sequences across time (stability) |
| 2.1.2. | Single-group model with YA_OA ($n_{YA–OA}$ = 320) | γLC_TP1_LC_TP2 | $r$ = 0.88; $\Delta\chi^2$(df = 1) = 66.93; $p$ < 0.001 | Multimodal locus coeruleus integrity is correlated across time (stability) |

| | | | | |
|---|---|---|---|---|
| 2.1.6 | Single-group model with YA–OA ($n_{YA–OA}$ = 320) | γSN_TP1_SN_TP2 | $r = 0.67$; $\Delta\chi^2(df = 1) = 47.71$ $p < 0.001$ | Multimodal substantia nigra–ventral tegmental area integrity is correlated across time (stability) |
| Result section: **Locus coeruleus and substantia nigra–ventral tegmental area are associated with different aspects of late-life memory performance** | | | | |
| 1.3.1. (covar) | Multi-group model with YA, OA ($n_{YA}$ = 69; $n_{OA}$ = 251) | γLC_WM'; γLC_EM'; γLC_Gf' | $\Delta\chi^2(df = 3) = 25.11$; $p < 0.001$ | Multimodal locus coeruleus integrity is associated with late-life cognition |
| 1.3.1. (covar) | Multi-group model with YA, OA ($n_{YA}$ = 69; $n_{OA}$ = 251) | γSN_WM'; γSN_EM'; γSN_Gf' | $\Delta\chi^2(df = 3) = 7.86$; $p = 0.049$ | Multimodal substantia nigra–ventral tegmental area integrity is associated with late-life cognition |
| 1.3.1. (covar) | Multi-group model with YA, OA ($n_{YA}$ = 69; $n_{OA}$ = 251) | γLC_SN' | $r = 0.25$; $\Delta\chi^2(df = 1) = 5.75$; $p = 0.017$ | Multimodal locus coeruleus and substantia nigra–ventral tegmental area integrity are correlated in older adults |
| 1.3.1. (covar) | Multi-group model with YA, OA ($n_{YA}$ = 69; $n_{OA}$ = 251) | γLC_WM'; γLC_EM'; γLC_Gf'; γSN_WM'; γSN_EM'; γSN_Gf' | $\Delta\chi^2(df = 3) = 15.66$; $p = 0.001$ | Multimodal locus coeruleus and substantia nigra–ventral tegmental area integrity are differentially associated with late-life cognition |
| 1.3.1. (covar) | Multi-group model with YA, OA ($n_{YA}$ = 69; $n_{OA}$ = 251) | γLC_EM' | $r = 0.49$; $\Delta\chi^2(df = 1) = 21.44$; $p < 0.001$ | Multimodal locus coeruleus integrity is associated with late-life episodic memory |
| 1.3.1. (covar) | Multi-group model with YA, OA ($n_{YA}$ = 69; $n_{OA}$ = 251) | γLC_EM'; γLC_WM'; γLC_Gf'; | $\Delta\chi^2(df = 2) = 10.64$; $p = 0.005$ | The association of multimodal locus coeruleus integrity with late-life episodic memory differs from the associations with working memory and fluid intelligence |
| 1.3.1. (covar) | Multi-group model with YA, OA ($n_{YA}$ = 69; $n_{OA}$ = 251) | γLC_EM'; γSN_EM'; | $\Delta\chi^2(df = 1) = 6.63$; $p = 0.01$ | The association of multimodal locus coeruleus integrity with late-life episodic memory differs from the association of multimodal substantia nigra–ventral tegmental area integrity with late-life episodic memory |
| 1.3.1. (covar) | Multi-group model with YA, OA ($n_{YA}$ = 69; $n_{OA}$ = 251) | γSN_WM' | $r = 0.28$; $\Delta\chi^2(df = 1) = 6.76$; $p = 0.009$ | Multimodal substantia nigra–ventral tegmental area integrity is associated with late-life working memory |
| 1.3.1. (covar) | Multi-group model with YA, OA ($n_{YA}$ = 69; $n_{OA}$ = 251) | γSN_WM'; γSN_EM'; γSN_Gf'; | $\Delta\chi^2(df = 2) = 5.73$; $p = 0.057$ | The association of multimodal substantia nigra–ventral tegmental area integrity with late-life working memory differs (on a trend level) from the associations with episodic memory and fluid intelligence |

| | | | | |
|---|---|---|---|---|
| 1.3.1. (covar) | Multi-group model with YA, OA ($n_{YA}$ = 69; $n_{OA}$ = 251) | γSN_WM'; γLC_WM'; | $\Delta\chi^2(df = 1) = 2.01$; $p = 0.156$ | The association of multimodal substantia nigra–ventral tegmental area integrity with late-life working memory differs from the association of multimodal locus coeruleus integrity with late-life working memory |
| 1.3.1. (reg) | Multi-group model with YA, OA ($n_{YA}$ = 69; $n_{OA}$ = 251) | γLC_EM' | $\beta = 0.5$; $\Delta\chi^2(df = 1) = 19.55$; $p < 0.001$ | Multimodal locus coeruleus integrity is associated with late-life episodic memory, even when accounting for substantia nigra–ventral tegmental area integrity |
| 1.3.1. (reg) | Multi-group model with YA, OA ($n_{YA}$ = 69; $n_{OA}$ = 251) | γSN_WM' | $\beta = 0.28$; $\Delta\chi^2(df = 1) = 6.05$; $p = 0.014$ | Multimodal substantia nigra–ventral tegmental area integrity is associated with late-life working memory, even when accounting for locus coeruleus integrity |

Result section:
**Locus coeruleus and substantia nigra–ventral tegmental area are associated with memory performance over and above medial temporal lobe volumes**

| | | | | |
|---|---|---|---|---|
| 1.3.2. (covar) | Single-group model with OA ($n_{OA}$ = 251) | γLC_MTL | $r = 0.41$; $\Delta\chi^2(df = 1) = 27.45$; $p < 0.001$ | Multimodal locus coeruleus integrity is associated with medial temporal lobe volume |
| 1.3.2. (covar) | Single-group model with OA ($n_{OA}$ = 251) | γSN_MTL | $r = 0.23$; $\Delta\chi^2(df = 1) = 6.29$; $p = 0.012$ | Multimodal substantia nigra–ventral tegmental area integrity is associated with medial temporal lobe volume |
| 1.3.2. (covar) | Single-group model with OA ($n_{OA}$ = 251) | γMTL_EM | $r = 0.33$; $\Delta\chi^2(df = 1) = 14.22$; $p < 0.001$ | Medial temporal lobe volume is associated with late-life episodic memory |
| 1.3.2. (reg) | Single-group model with OA ($n_{OA}$ = 251) | γLC_EM | $\beta = 0.43$; $\Delta\chi^2(df = 1) = 11.96$; $p < 0.001$ | Multimodal locus coeruleus integrity is associated with late-life episodic memory, even when accounting for medial temporal lobe volume and multimodal substantia nigra–ventral tegmental area integrity |
| 1.3.2. (reg) | Single-group model with OA ($n_{OA}$ = 251) | γMTL_EM | $\beta = 0.16$; $\Delta\chi^2(df = 1) = 2.46$; $p = 0.117$ | Medial temporal lobe volume is associated with late-life episodic memory, when accounting for the integrity of catecholaminergic nuclei |
| 1.3.2. (reg) | Single-group model with OA ($n_{OA}$ = 251) | γSN_WM | $\beta = 0.28$; $\Delta\chi^2(df = 1) = 5.8$; $p = 0.016$ | Multimodal substantia nigra–ventral tegmental area integrity is associated with late-life working memory, even when accounting for medial temporal lobe volume and multimodal locus coeruleus integrity |

Result section:
**Longitudinal changes in locus coeruleus integrity predict future episodic memory performance**

| | | | | |
|---|---|---|---|---|
| 2.1.3. | Single-group model with OA ($n_{OA}$ = 251) | γLCmt_slope_LCnomt_slope | $r = 0.16$; $\Delta\chi^2(df = 1) = 6.09$; $p = 0.014$ | Late-life changes in locus coeruleus intensity are correlated across MRI sequences |

| | | | | |
|---|---|---|---|---|
| 2.1.7. | Single-group model with OA ($n_{OA}$ = 251) | γSNmt_slope_SNnomt_slope | $r$ = 0.13; $\Delta\chi^2(df = 1)$ = 5.91; $p$ = 0.015 | Late-life changes in substantia nigra–ventral tegmental area intensity are correlated across MRI sequences |
| 2.1.4. | Single-group model with OA ($n_{OA}$ = 251) | σLC_slope | $\Delta\chi^2(df = 1)$ = 6.09; $p$ = 0.014 | There are reliable individual differences in multimodal locus coeruleus integrity change |
| 2.1.8. | Single-group model with OA ($n_{OA}$ = 251) | σSN_slope | $\Delta\chi^2(df = 1)$ = 5.91; $p$ = 0.015 | There are reliable individual differences in multimodal substantia nigra–ventral tegmental area integrity change |
| 2.3.1. | Single-group model with OA ($n_{OA}$ = 251) | γAge_TP2_LC_slope | $\beta$ = –0.18; $\Delta\chi^2(df = 1)$ = 4.81; $p$ = 0.028 | Chronological age is associated with change in multimodal locus coeruleus integrity |
| 2.3.2. | Single-group model with OA ($n_{OA}$ = 251) | γAge_TP2_SN_slope | $\beta$ = –0.29; $\Delta\chi^2(df = 1)$ = 3.95; $p$ = 0.047 | Chronological age is associated with change in multimodal substantia nigra–ventral tegmental area integrity |
| 2.3.1. | Single-group model with OA ($n_{OA}$ = 251) | γLC_slope_EM_TP3 | $\beta$ = 0.23; $\Delta\chi^2(df = 1)$ = 4.73; $p$ = 0.03 | Change in multimodal locus coeruleus integrity is associated with subsequent episodic memory |
| 2.3.2. | Single-group model with OA ($n_{OA}$ = 251) | γSN_slope_WM_TP3 | $\beta$ = 0.27; $\Delta\chi^2(df = 1)$ = 1.55; $p$ = 0.213 | Change in multimodal substantia nigra–ventral tegmental area integrity is not significantly associated with subsequent working memory |

**Note**: μ, mean; σ, variance; γ, covariance or regression; Path names including “ ' ” indicate a multi-group model with age group-specific parameter estimates; covar, covariance model; reg, multiple regression model; All structural equation models were estimated with full-information maximum likelihood estimation (that is, cases with partially missing data were not excluded [6]).

Cross-sectional neural models:

**Table S2.** Model fit and invariance for cross-sectional neural models

| Model number | Model name | Age group | Time point | Invariance | $\chi^2$ | *df* | *p* | RMSEA | CFI |
|---|---|---|---|---|---|---|---|---|---|
| 1.1.1 | LC and SN–VTA modality-specific factors | YA, OA | 2 | Strict (across age groups) | 111.131 | 82 | 0.018 | 0.047 | 0.963 |
| 1.1.2 | LC and SN–VTA multimodal factors | YA, OA | 2 | Partial strict[1] (across age groups) | 109.891 | 88 | 0.057 | 0.039 | 0.972 |
| 1.1.3 | MTL regional factors | OA | 2 | – (single-group, cross-sectional model) | 55.409 | 1 | < 0.001 | 0.467[2] | 0.952 |

*Note:* LC, locus coeruleus; SN–VTA, substantia nigra–ventral tegmental area; MTL, medial temporal lobe; YA, younger adults; OA, older adults; YA, OA, multi-group model including both age groups; RMSEA, root mean square error of approximation; CFI, comparative fit index

[1] Partial strict invariance for model 1.1.2—the model did not show age-invariant intercepts of the modality-specific LC factors (test for strong invariance). We thus allowed for differences in µLCmt– across groups (i.e., partial invariance). Note that this does not influence our following analyses, which are based on covariances and regressions (not intercepts).

[2] Model 1.1.3 exceeds conventional recommendations for RMSEA. Note, however, that the unified neuro–cognitive model (1.3.2) on which we base our inferences shows good fit [7,8].

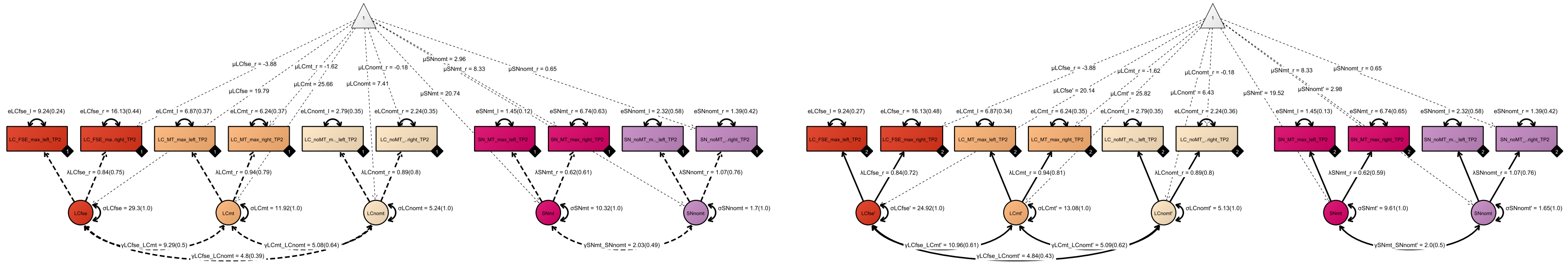

**Figure S3**. Model 1.1.1.

Pictorial rendition of a confirmatory factor analysis including modality-specific LC and SN–VTA factors for younger and older adults. Rectangles and circles indicate manifest (observed) and latent variables, respectively. The constant is depicted by a triangle. Black diamonds on manifest variables indicate the age group. The younger adult submodel is represented by dashed lines (◆1), and the older adults submodel is represented by solid lines (◆2). (Co)Variances (γ, σ) and loadings (λ) in brackets indicate standardized estimates. Parameters that have the same name are constrained to be equal across age groups. One-headed arrows indicate regressions, double-headed arrows indicate correlations.

LC, locus coeruleus; SN, substantia nigra–ventral tegmental area; fse, Fast Spin Echo; mt, Magnetization Transfer (MT+)

nomt, Proton Density (MT–)

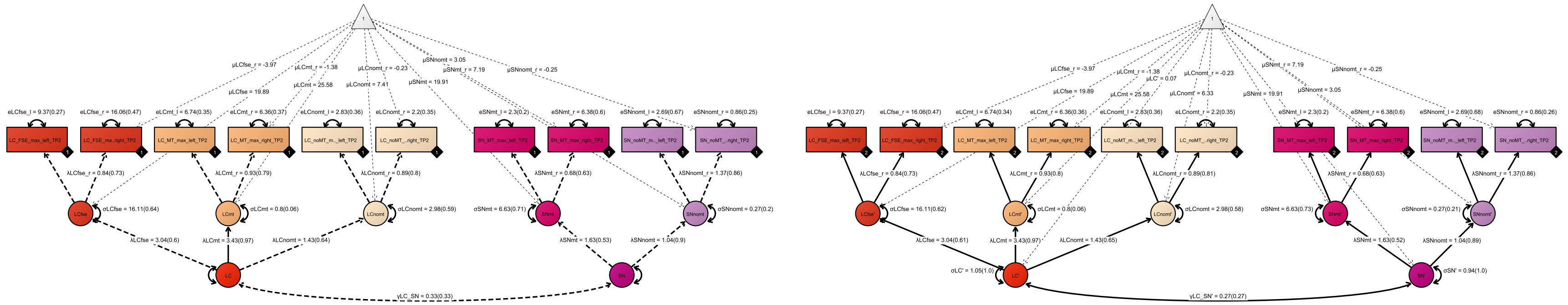

**Figure S4**. Model 1.1.2.

Pictorial rendition of a confirmatory factor analysis including multimodal LC and SN–VTA factors for younger and older adults. Rectangles and circles indicate manifest (observed) and latent variables, respectively. The constant is depicted by a triangle. Black diamonds on manifest variables indicate the age group. The younger adult submodel is represented by dashed lines (◆1), and the older adults submodel is represented by solid lines (◆2). (Co)Variances (γ, σ) and loadings (λ) in brackets indicate standardized estimates. Parameters that have the same name are constrained to be equal across age groups. One-headed arrows indicate regressions, double-headed arrows indicate correlations.
LC, locus coeruleus; SN, substantia nigra–ventral tegmental area; fse, Fast Spin Echo; mt, Magnetization Transfer (MT+); nomt, Proton Density (MT–).

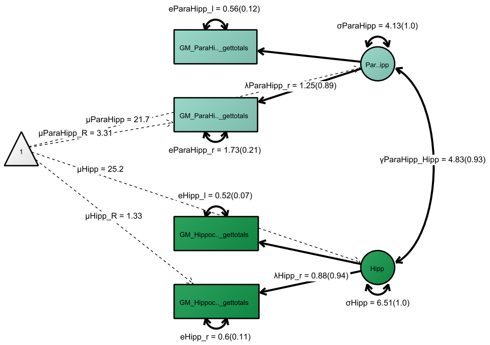

**Figure S5**. Model 1.1.3.

Pictorial rendition of a confirmatory factor analysis including regional factors for hippocampal and parahippocampal volume for older adults. Rectangles and circles indicate manifest (observed) and latent variables, respectively. The constant is depicted by a triangle. (Co)Variances (γ, σ) and loadings (λ) in brackets indicate standardized estimates. One-headed arrows indicate regressions, double-headed arrows indicate correlations. Hipp, hippocampus; Parahipp, parahippocampal cortex.

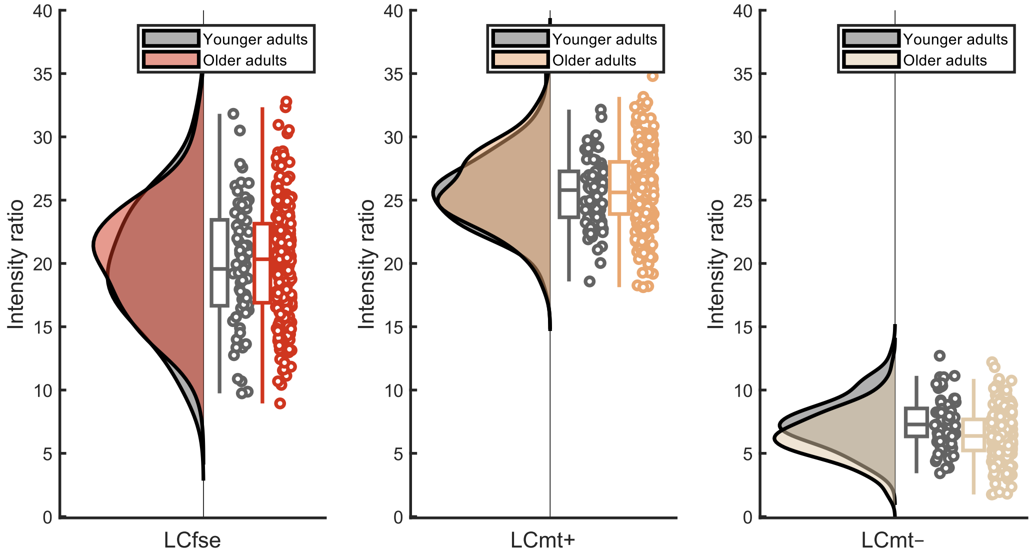

**Figure S6**. Cross-sectional age differences in modality-specific LC factors.

Visualized data are based on the statistical model 1.1.1.
Raincloud plots based on [9]. LC, locus coeruleus; fse, Fast Spin Echo; mt, Magnetization Transfer (MT+); nomt, Proton Density (MT–).

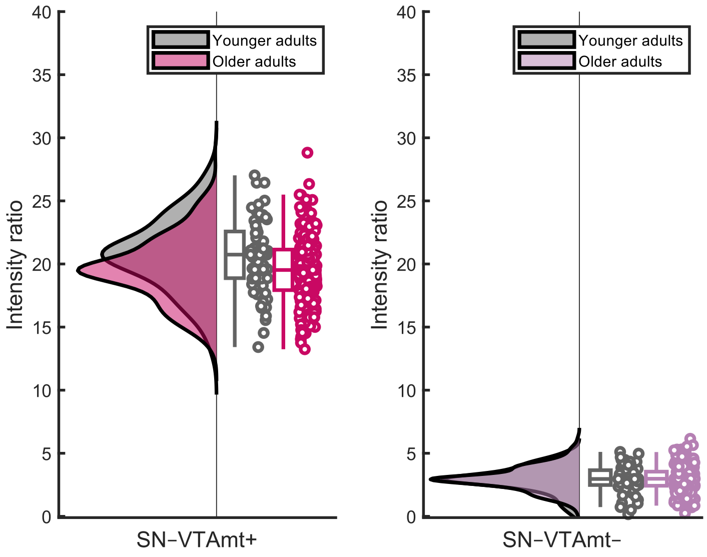

**Figure S7.** Cross-sectional age differences in modality-specific SN–VTA factors.

Visualized data are based on the statistical model 1.1.1.
Raincloud plots based on [9]. SN–VTA, substantia nigra–ventral tegmental area; fse, Fast Spin Echo; mt, Magnetization Transfer (MT+); nomt, Proton Density (MT–).

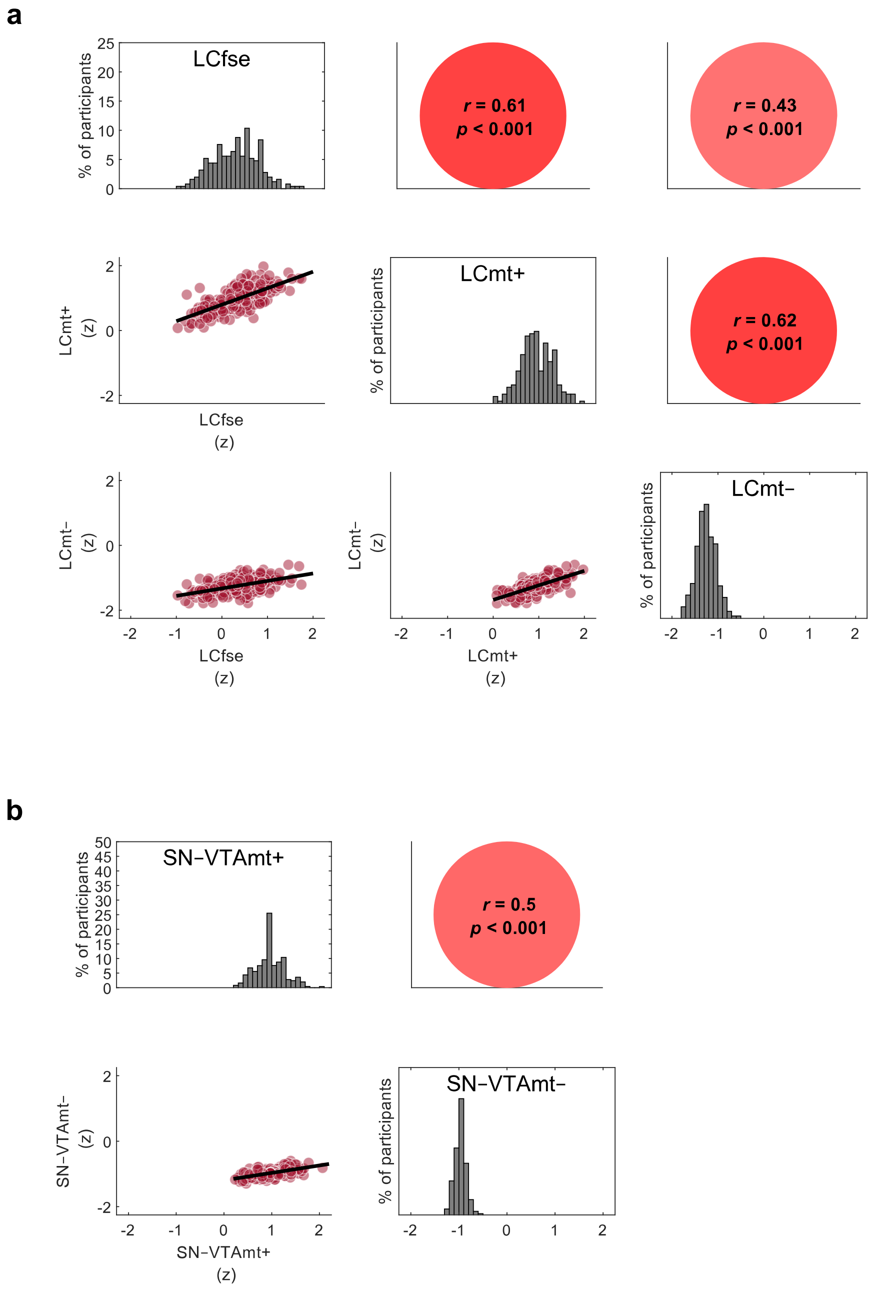

**Figure S8.** LC and SN–VTA intensities are correlated across imaging modalities (in older adults).

Visualized data are based on model 1.1.1. Note, the diagonal shows intensity, standardized across all sequences, to facilitate comparing intensity distributions. Imaging sequences included a Fast Spin Echo (FSE) sequence, and a Magnetization Transfer sequence, acquired once with a dedicated magnetic saturation pulse (MT+) and once without, yielding a proton density image (MT–). LC, locus coeruleus; SN–VTA, substantia nigra–ventral tegmental area.

Cross-sectional cognitive models:

**Table S3**. Model fit and invariance for cross-sectional cognitive models

| Model number | Model name | Age group | Time point | Invariance | $\chi^2$ | *df* | *p* | RMSEA | CFI |
|---|---|---|---|---|---|---|---|---|---|
| 1.2.1 | WM, EM, and Gf factors | YA, OA | 2 | Strong (across age groups) | 104.934 | 78 | 0.023 | 0.047 | 0.966 |

*Note:* WM, working memory; EM, episodic memory; Gf, fluid intelligence; YA, younger adults; OA, older adults; YA, OA, multi-group model including both age groups; RMSEA, root mean square error of approximation; CFI, comparative fit index

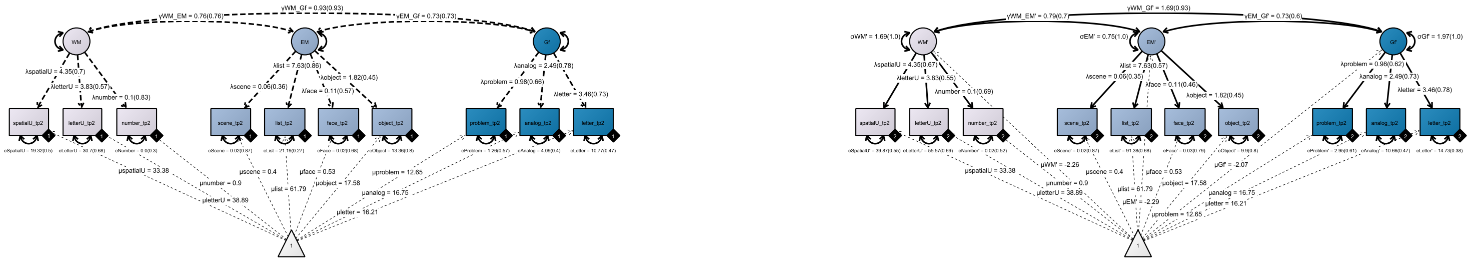

**Figure S9.** Model 1.2.1.

Pictorial rendition of a confirmatory factor analysis including cognitive factors for working memory, episodic memory, and fluid intelligence for younger and older adults.

Rectangles and circles indicate manifest (observed) and latent variables, respectively. The constant is depicted by a triangle. Black diamonds on manifest variables indicate the age group. The younger adult submodel is represented by dashed lines (◆1), and the older adults submodel is represented by solid lines (◆2). (Co)Variances (γ, σ) and loadings (λ) in brackets indicate standardized estimates. Parameters that have the same name are constrained to be equal across age groups. One-headed arrows indicate regressions, double-headed arrows indicate correlations.

WM, working memory; EM, episodic memory; Gf, fluid intelligence.

Cross-sectional neuro–cognitive models:

**Table S4**. Model fit and invariance for cross-sectional neuro–cognitive models

| Model number | Model name | Age group | Time point | Invariance | $\chi^2$ | *df* | *p* | RMSEA | CFI |
|---|---|---|---|---|---|---|---|---|---|
| 1.3.1 | Covariances or regressions[1] between: LC, SN–VTA and WM, EM, Gf factors | YA, OA | 2 | – (invariance tested for cognitive and neural submodels) | 463.293 | 354 | < 0.001 | 0.044 | 0.934 |
| 1.3.2 | Covariances or regressions between: MTL, LC, SN–VTA and WM, EM, Gf factors | OA | 2 | – (single-group, cross-sectional model) | 364.132 | 230 | < 0.001 | 0.048 | 0.944 |

*Note:* LC, locus coeruleus; SN–VTA, substantia nigra–ventral tegmental area; MTL, medial temporal lobe; WM, working memory; EM, episodic memory; Gf, fluid intelligence; YA, younger adults; OA, older adults; YA, OA, multi-group model including both age groups; RMSEA, root mean square error of approximation; CFI, comparative fit index

[1] The models using latent covariances and regressions are statistically equivalent. Thus, we provide the invariance and fit for them together.

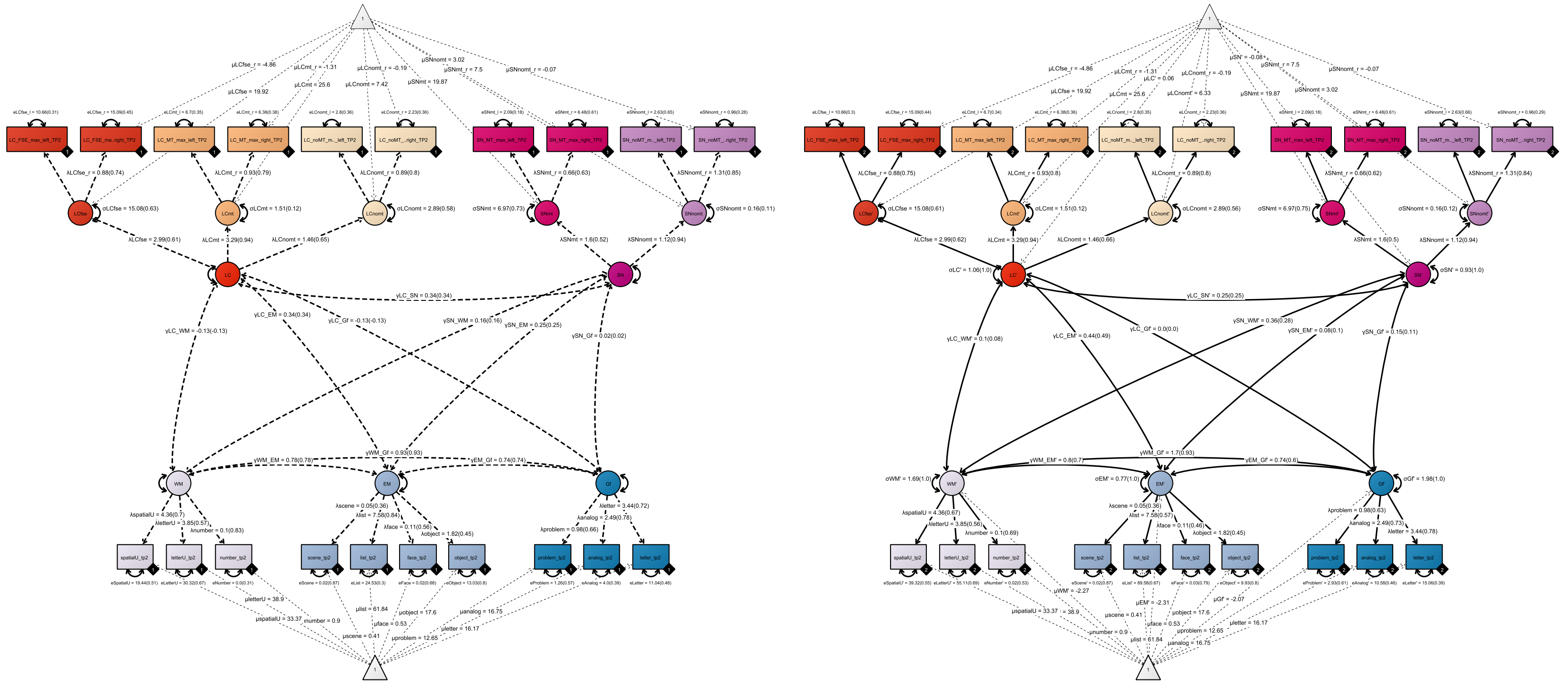

**Figure S10. Model 1.3.1. (a)**

Pictorial rendition of a structural equation model probing the association (correlation) between LC and SN–VTA integrity and cognitive factors for working memory, episodic memory, and fluid intelligence in younger and older adults.
Rectangles and circles indicate manifest (observed) and latent variables, respectively. The constant is depicted by a triangle. Black diamonds on manifest variables indicate the age group. The younger adult submodel is represented by dashed lines (◆1), and the older adults submodel is represented by solid lines (◆2). (Co)Variances (γ, σ) and loadings (λ) in brackets indicate standardized estimates. Parameters that have the same name are constrained to be equal across age groups. One-headed arrows indicate regressions, double-headed arrows indicate correlations.
LC, locus coeruleus; SN, substantia nigra–ventral tegmental area; fse, Fast Spin Echo; mt, Magnetization Transfer (MT+); nomt, Proton Density (MT–); WM, working memory; EM, episodic memory; Gf, fluid intelligence.

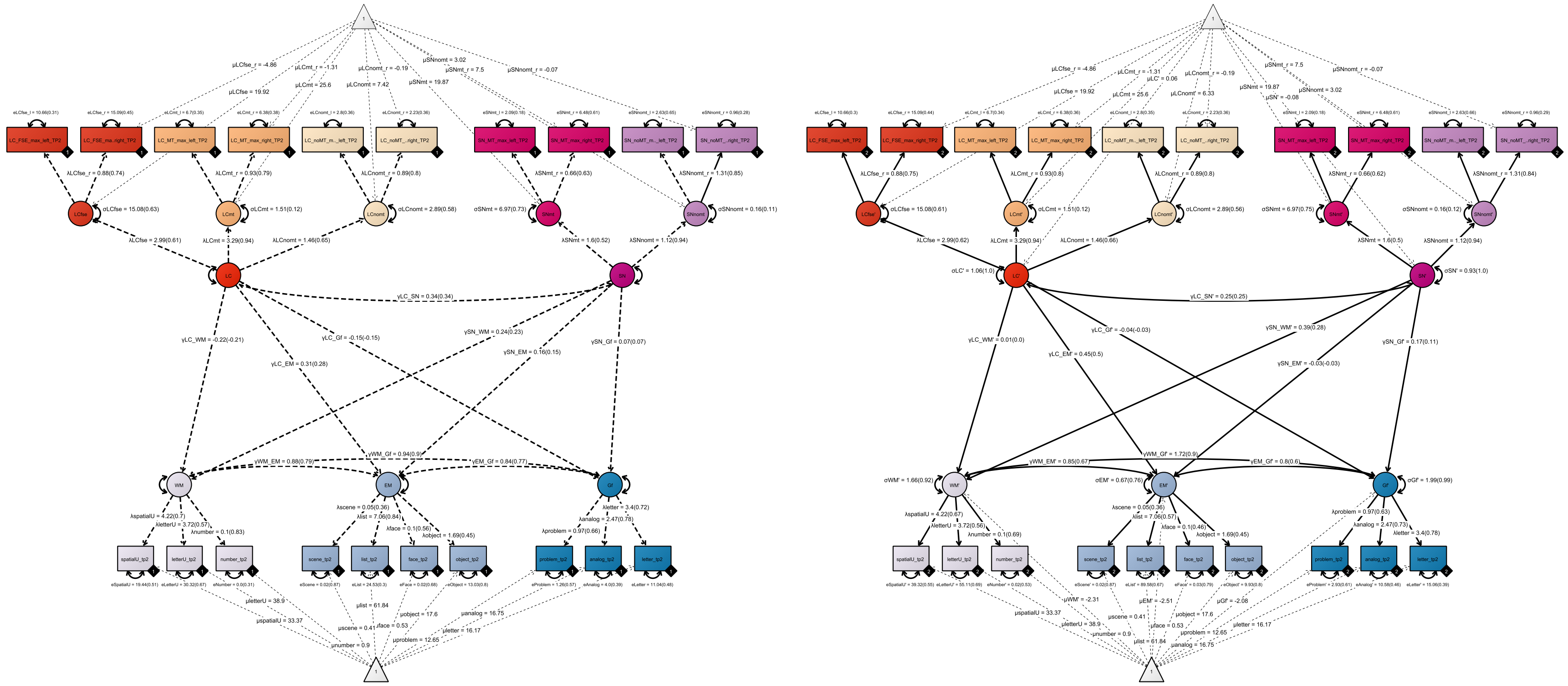

**Figure S11. Model 1.3.1. (b)**

Pictorial rendition of a structural equation model probing unique associations (multiple regression) between LC and SN–VTA integrity and cognitive factors for working memory, episodic memory, and fluid intelligence in younger and older adults.
Rectangles and circles indicate manifest (observed) and latent variables, respectively. The constant is depicted by a triangle. Black diamonds on manifest variables indicate the age group. The younger adult submodel is represented by dashed lines (◆1), and the older adults submodel is represented by solid lines (◆2). (Co)Variances (γ, σ) and loadings (λ) in brackets indicate standardized estimates. Parameters that have the same name are constrained to be equal across age groups. One-headed arrows indicate regressions, double-headed arrows indicate correlations.
LC, locus coeruleus; SN, substantia nigra–ventral tegmental area; fse, Fast Spin Echo; mt, Magnetization Transfer (MT+); nomt, Proton Density (MT–); WM, working memory; EM, episodic memory; Gf, fluid intelligence.

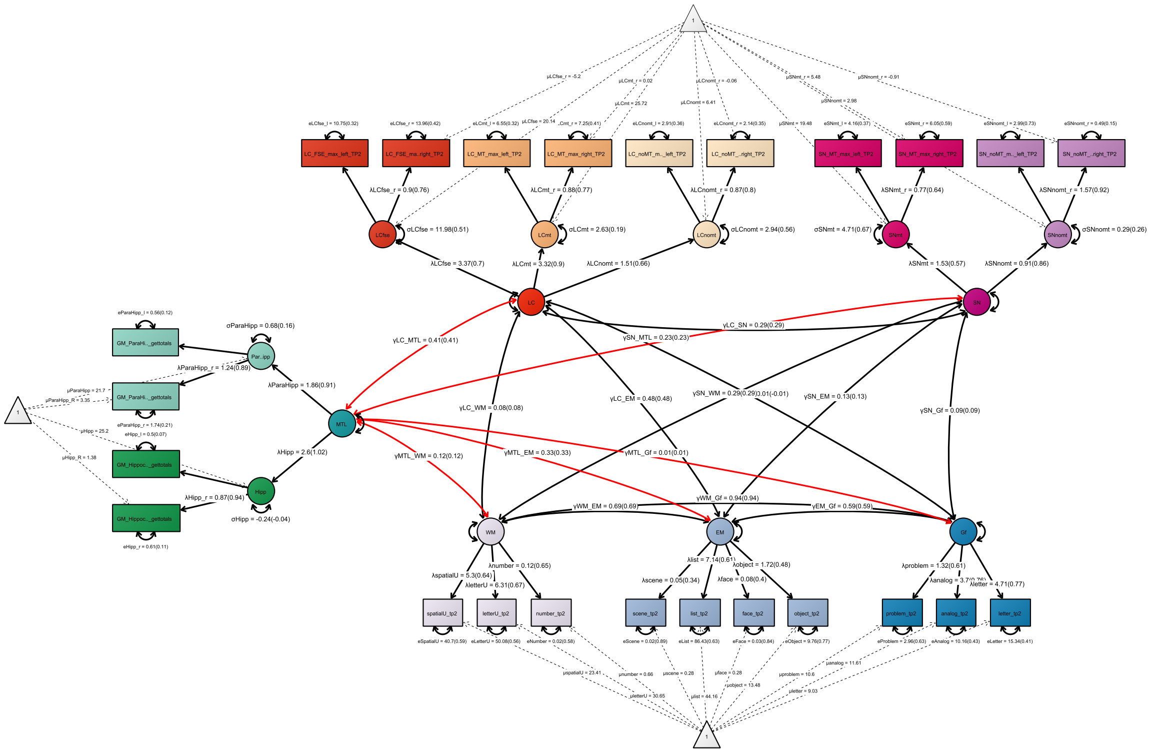

**Figure S12. Model 1.3.2. (a)**

Pictorial rendition of a structural equation model probing the association (correlation) between LC and SN–VTA integrity, medial temporal lobe volume and cognitive factors for working memory, episodic memory, and fluid intelligence in older adults.
Rectangles and circles indicate manifest (observed) and latent variables, respectively. The constant is depicted by a triangle. (Co)Variances (γ, σ) and loadings (λ) in brackets indicate standardized estimates. One-headed arrows indicate regressions, double-headed arrows indicate correlations.
LC, locus coeruleus; SN, substantia nigra–ventral tegmental area; fse, Fast Spin Echo; mt, Magnetization Transfer (MT+); nomt, Proton Density (MT–); Hipp, hippocampus; Parahipp, parahippocampal cortex; WM, working memory; EM, episodic memory; Gf, fluid intelligence.

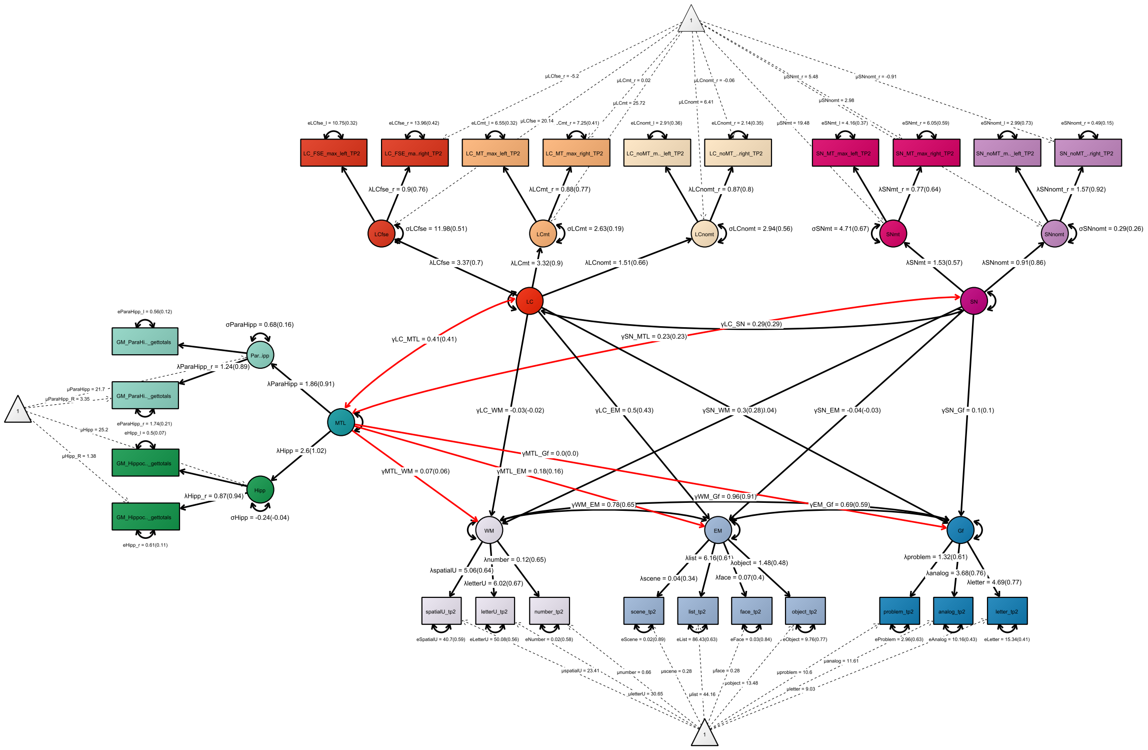

**Figure S13. Model 1.3.2. (b)**

Pictorial rendition of a structural equation model probing unique associations (multiple regression) between LC and SN–VTA integrity, medial temporal lobe volume and cognitive factors for working memory, episodic memory, and fluid intelligence in older adults.
Rectangles and circles indicate manifest (observed) and latent variables, respectively. The constant is depicted by a triangle. (Co)Variances (γ, σ) and loadings (λ) in brackets indicate standardized estimates. One-headed arrows indicate regressions, double-headed arrows indicate correlations.
LC, locus coeruleus; SN, substantia nigra–ventral tegmental area; fse, Fast Spin Echo; mt, Magnetization Transfer (MT+); nomt, Proton Density (MT–); Hipp, hippocampus; Parahipp, parahippocampal cortex; WM, working memory; EM, episodic memory; Gf, fluid intelligence.

### Cross-sectional neuro–cognitive model with mean intensity ratios

Control analyses using mean intensity ratios largely recapitulate our main findings. That is, we fit model 1.32 b using mean intensity ratios instead of peak intensity ratios for each catecholaminergic nuclei (model fit: CFI = 0.939; RMSEA = 0.054. Heywood case for σHipp and σSNnomt). In this control analysis, we again find an (A) interrelation of intensity ratios across MRI modalities; (B) positive coupling of multimodal locus coeruleus and substantia nigra–ventral tegmental area factors; (C) association of neuromodulatory integrity factors and medial-temporal lobe volumes; (D) association of locus coeruleus integrity and episodic memory performance. However, numerical comparisons of the magnitude of brain–cognition associations across analysis approaches (peak vs mean intensity) suggests that the peak intensity metric better isolates behaviorally-relevant hyperintensities within the search spaces. Due to the Heywood cases, the mean intensity model should be interpreted with caution. We provide statistical comparisons of the model parameters below.

**Table S5**. Comparison of parameter estimates of cross-sectional neuro-cognitive models fit with peak and mean intensity ratio data

| | **Standardized estimate** | | **Test for difference of coefficients** | | |
|---|---|---|---|---|---|
| **Path** | **Mean ratios** | **Peak ratios** | ***df*** | **$\Delta\chi^2$** | ***p*** |
| γLC_SN | 0.43 | 0.29 | 1 | 2.03 | 0.154 |
| γLC_MTL | 0.42 | 0.41 | 1 | 0.01 | 0.920 |
| γSN_MTL | 0.34 | 0.23 | 1 | 1.28 | 0.258 |
| γLC_EM | 0.22 | 0.43 | 1 | 3.46 | 0.063 |
| γSN_WM | –0.03 | 0.28 | 1 | 6.93 | 0.009 |

Longitudinal neural models:

**Table S6**. Model fit and invariance for longitudinal neural models

| Model number | Model name | Age group | Time point | Invariance | $\chi^2$ | *df* | *p* | RMSEA | CFI |
|---|---|---|---|---|---|---|---|---|---|
| 2.1.1 | Covariance of modality-specific LC factors over time | YA–OA | 1, 2 | Strong (across time points) | 53.468 | 25 | 0.001 | 0.06 | 0.97 |
| 2.1.2 | Covariance of multimodal LC factors over time | YA–OA | 1, 2 | Weak[1] (across time points) | 23.427 | 14 | 0.0537 | 0.046 | 0.987 |
| 2.1.3 | Covariance of modality-specific LC change factors | OA | 1, 2 | – (single-group, latent-change score models) | 1.288 | 2 | 0.525 | ~ 0 | ~ 1 |
| 2.1.4 | Multimodal LC change factor | OA | 1, 2 | – (single-group, latent-change score model) | 1.288 | 1 | 0.256 | 0.034 | 0.998 |
| 2.1.5 | Covariance of modality-specific SN–VTA factors over time | YA–OA | 1, 2 | Strict (across time points) | 25.167 | 18 | 0.120 | 0.035 | 0.982 |
| 2.1.6 | Covariance of multimodal SN–VTA factors over time | YA–OA | 1, 2 | Strict (across time points) | 36.589 | 23 | 0.036 | 0.043 | 0.966 |
| 2.1.7 | Covariance of modality-specific SN–VTA change factors | OA | 1, 2 | – (single-group, latent-change score models) | 4.866 | 2 | 0.088 | 0.076 | 0.96 |
| 2.1.8 | Multimodal SN–VTA change factor | OA | 1, 2 | – (single-group, latent-change score model) | 4.866 | 1 | 0.027 | 0.124[2] | 0.947 |

*Note:* LC, locus coeruleus; SN–VTA, substantia nigra–ventral tegmental area; YA, younger adults; OA, older adults; YA–OA, single group model including both age groups; RMSEA, root mean square error of approximation; CFI, comparative fit index.

[1] Model 2.1.2 does not show invariant modality-specific LC intercepts over time. Thus, strong invariance constraints do not hold. However, as we analyze covariances over time (not means), this does not influence any of the reported findings.

[2] Model 2.1.7 exceeds conventional recommendations for RMSEA. Note, however, that the unified neuro–cognitive model (2.3.2) on which we base our inferences shows good fit [8,10].

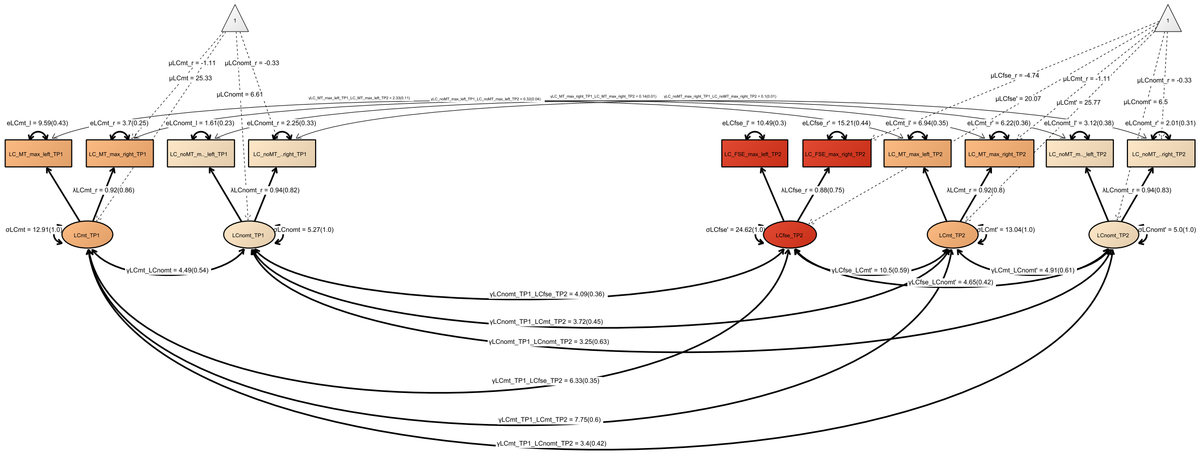

**Figure S14.** Model 2.1.1.

Pictorial rendition of a confirmatory factor analysis including modality-specific LC factors for time point 1 and 2 across younger and older adults. Rectangles and circles indicate manifest (observed) and latent variables, respectively. The constant is depicted by a triangle. (Co)Variances (γ, σ) and loadings (λ) in brackets indicate standardized estimates. One-headed arrows indicate regressions, double-headed arrows indicate correlations. LC, locus coeruleus; SN, substantia nigra–ventral tegmental area; fse, Fast Spin Echo; mt, Magnetization Transfer (MT+); nomt, Proton Density (MT–).

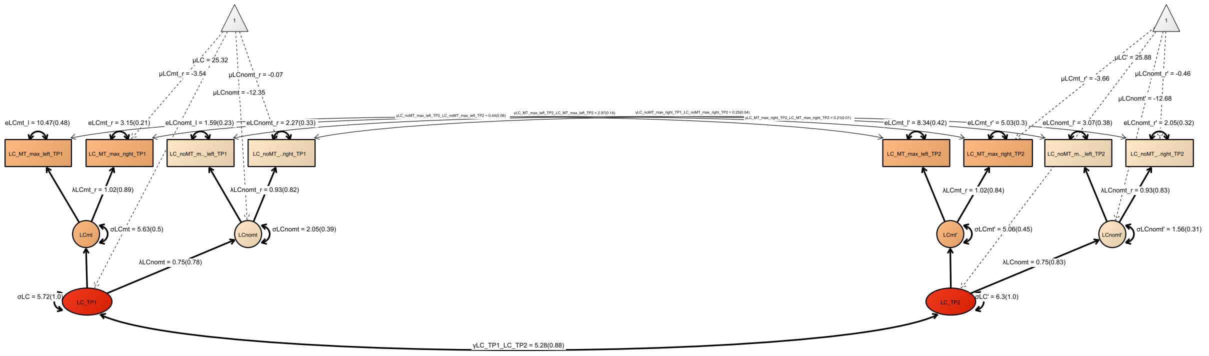

**Figure S15.** Model 2.1.2.

Pictorial rendition of a confirmatory factor analysis including multimodal LC factors for time point 1 and 2 across younger and older adults. Rectangles and circles indicate manifest (observed) and latent variables, respectively. The constant is depicted by a triangle. (Co)Variances (γ, σ) and loadings (λ) in brackets indicate standardized estimates. One-headed arrows indicate regressions, double-headed arrows indicate correlations. LC, locus coeruleus; SN, substantia nigra–ventral tegmental area; fse, Fast Spin Echo; mt, Magnetization Transfer (MT+); nomt, Proton Density (MT–); TP, time point.

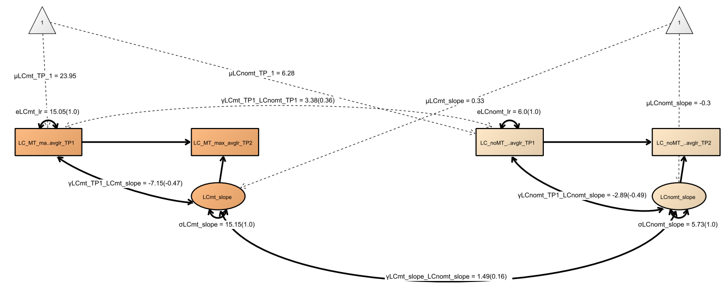

**Figure S16.** Model 2.1.3.

Pictorial rendition of modality-specific LC latent change score models including data of time point 1 and 2 of older adults. Rectangles and circles indicate manifest (observed) and latent variables, respectively. The constant is depicted by a triangle. (Co)Variances (γ, σ) and loadings (λ) in brackets indicate standardized estimates. One-headed arrows indicate regressions, double-headed arrows indicate correlations. LC, locus coeruleus; SN, substantia nigra–ventral tegmental area; fse, Fast Spin Echo; mt, Magnetization Transfer (MT+); nomt, Proton Density (MT–); TP, time point.

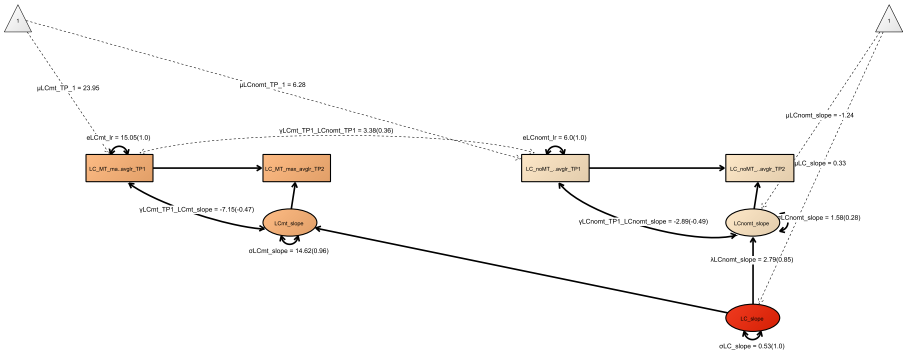

**Figure S17.** Model 2.1.4.

Pictorial rendition of a confirmatory factor analysis aggregating across modality-specific LC latent change score models (time point 1→2) in older adults. Rectangles and circles indicate manifest (observed) and latent variables, respectively. The constant is depicted by a triangle. (Co)Variances (γ, σ) and loadings (λ) in brackets indicate standardized estimates. One-headed arrows indicate regressions, double-headed arrows indicate correlations. LC, locus coeruleus; SN, substantia nigra–ventral tegmental area; fse, Fast Spin Echo; mt, Magnetization Transfer (MT+); nomt, Proton Density (MT–); TP, time point.

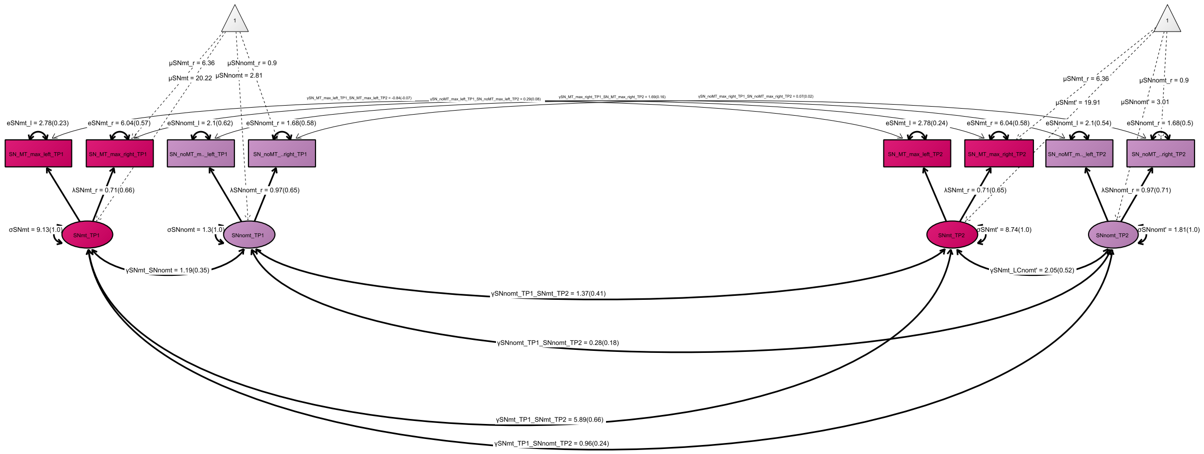

**Figure S18.** Model 2.1.5.

Pictorial rendition of a confirmatory factor analysis including modality-specific SN–VTA factors for time point 1 and 2 across younger and older adults. Rectangles and circles indicate manifest (observed) and latent variables, respectively. The constant is depicted by a triangle. (Co)Variances (γ, σ) and loadings (λ) in brackets indicate standardized estimates. One-headed arrows indicate regressions, double-headed arrows indicate correlations.
LC, locus coeruleus; SN, substantia nigra–ventral tegmental area; fse, Fast Spin Echo; mt, Magnetization Transfer (MT+); nomt, Proton Density (MT–); TP, time point.

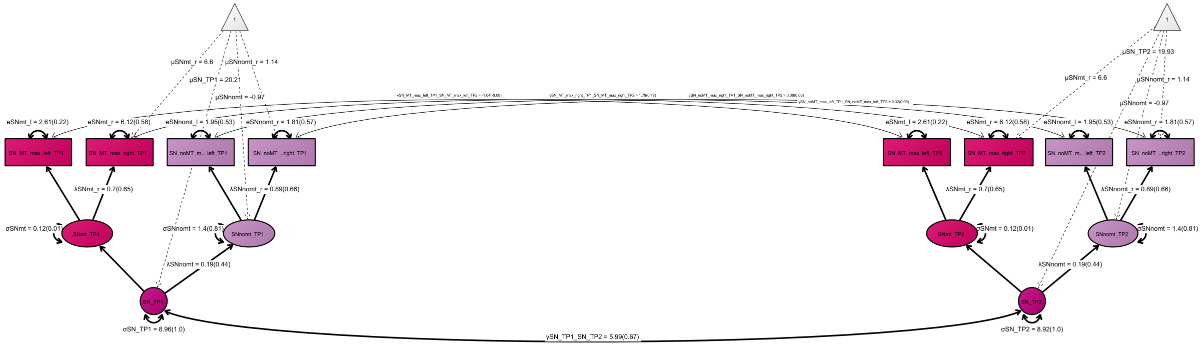

**Figure S19.** Model 2.1.6.

Pictorial rendition of a confirmatory factor analysis including multimodal SN–VTA factors for time point 1 and 2 across younger and older adults. Rectangles and circles indicate manifest (observed) and latent variables, respectively. The constant is depicted by a triangle. (Co)Variances (γ, σ) and loadings (λ) in brackets indicate standardized estimates. One-headed arrows indicate regressions, double-headed arrows indicate correlations. LC, locus coeruleus; SN, substantia nigra–ventral tegmental area; fse, Fast Spin Echo; mt, Magnetization Transfer (MT+); nomt, Proton Density (MT–); TP, time point.

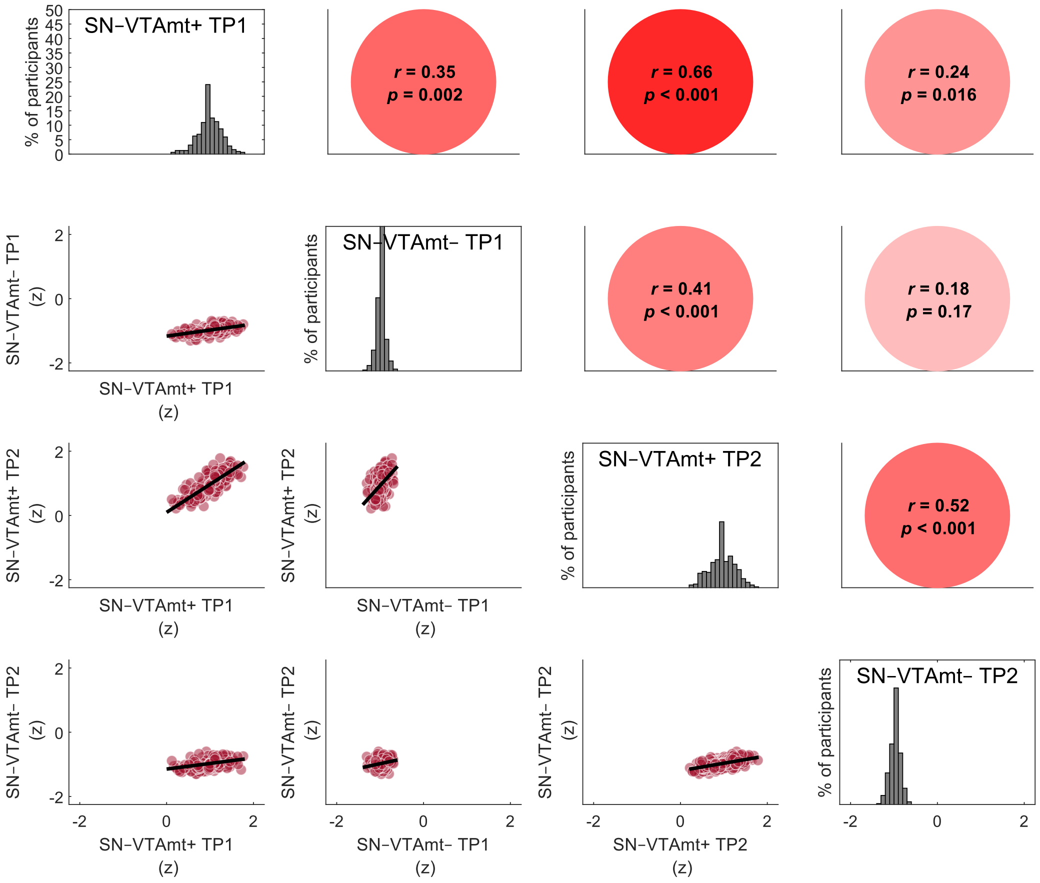

**Figure S20.** SN–VTA intensities are correlated across imaging modalities—a marker for their agreement—and time points—a marker for their stability (across younger and older adults).

Visualized data are based on the statistical model 2.1.5. For the same analyses using LC area data, see Figure 3. Note, the diagonal shows SN–VTA intensity, standardized across all sequences and time points, to facilitate comparing intensity distributions. Imaging sequences included a Magnetization Transfer sequence, acquired once with a dedicated magnetic saturation pulse (MT+) and once without, resulting in a proton density image (MT–). LC, locus coeruleus; SN–VTA, substantia nigra–ventral tegmental area; TP, time point.

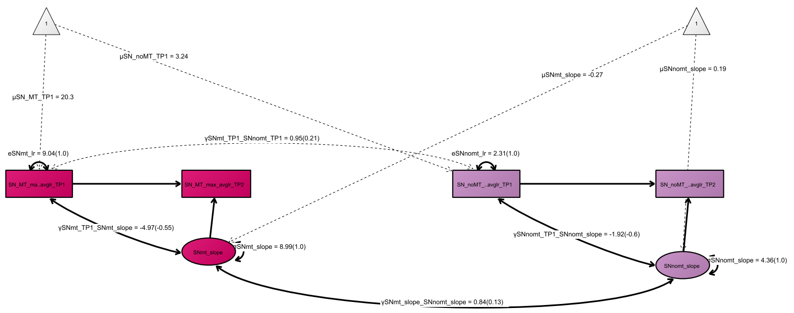

**Figure S21.** Model 2.1.7.

Pictorial rendition of modality-specific SN–VTA latent change scores models including data of time point 1 and 2 of older adults. Rectangles and circles indicate manifest (observed) and latent variables, respectively. The constant is depicted by a triangle. (Co)Variances (γ, σ) and loadings (λ) in brackets indicate standardized estimates. One-headed arrows indicate regressions, double-headed arrows indicate correlations. LC, locus coeruleus; SN, substantia nigra–ventral tegmental area; fse, Fast Spin Echo; mt, Magnetization Transfer (MT+); nomt, Proton Density (MT–); TP, time point.

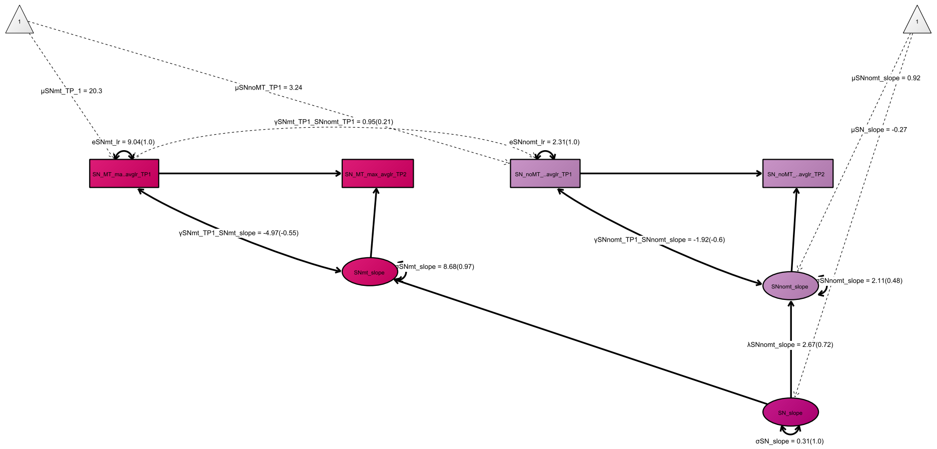

**Figure S22.** Model 2.1.8.

Pictorial rendition of a confirmatory factor analysis aggregating across modality-specific SN–VTA latent change score models (time point 1→2) in older adults. Rectangles and circles indicate manifest (observed) and latent variables, respectively. The constant is depicted by a triangle. (Co)Variances (γ, σ) and loadings (λ) in brackets indicate standardized estimates. One-headed arrows indicate regressions, double-headed arrows indicate correlations.
LC, locus coeruleus; SN, substantia nigra–ventral tegmental area; fse, Fast Spin Echo; mt, Magnetization Transfer (MT+); nomt, Proton Density (MT–); TP, time point.

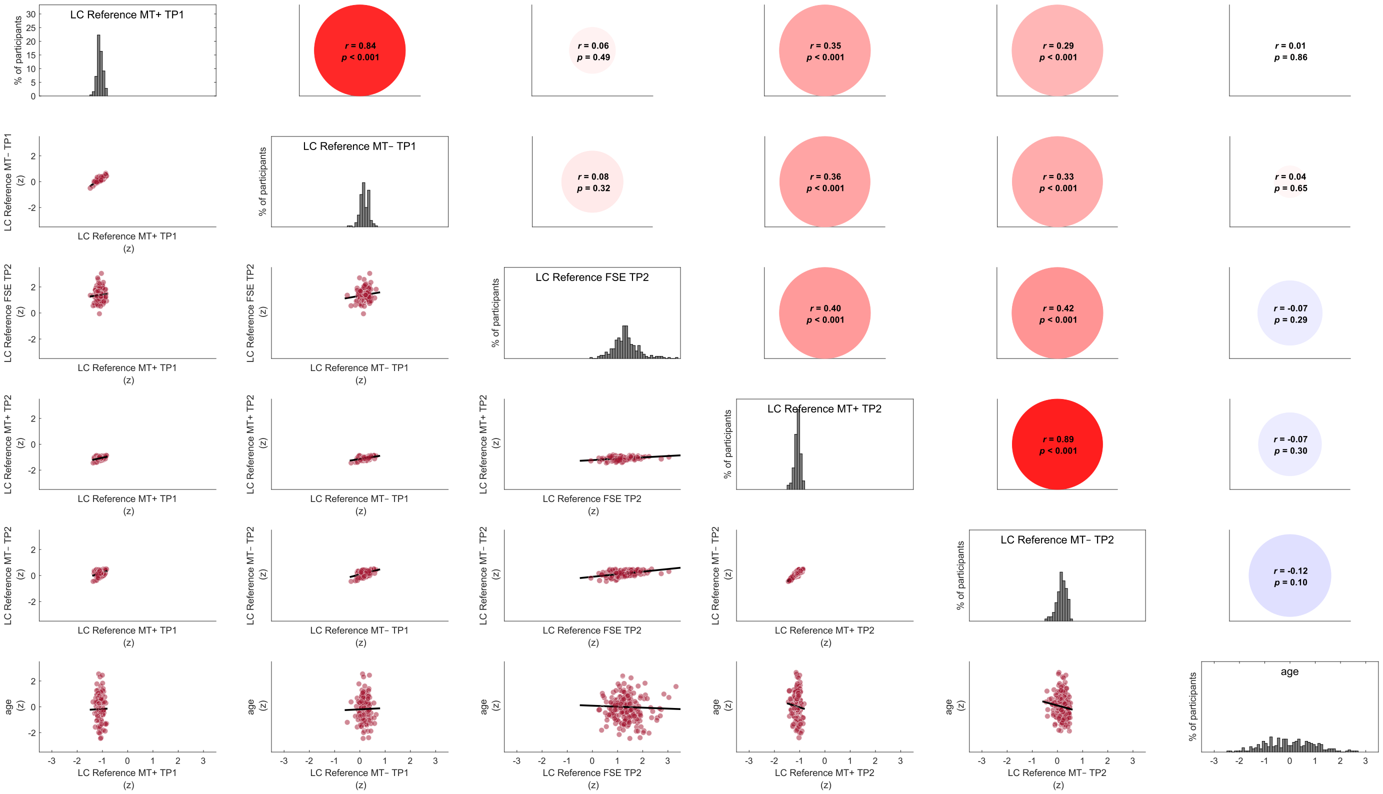

**Figure S23**. Pontine reference intensities across imaging modalities and time points, and their association with chronological age (in older adults). Visualized data are averaged across hemispheres. For the same analyses using LC data, see Figure 3. Note, the diagonal shows intensity, standardized across all sequences and time points, to facilitate comparing intensity distributions (age was standardized separately). Imaging sequences included a Fast Spin Echo sequence and a Magnetization Transfer sequence, acquired once with a dedicated magnetic saturation pulse (MT+) and once without, resulting in a proton density image (MT–). LC, locus coeruleus; SN–VTA, substantia nigra–ventral tegmental area; TP, time point.

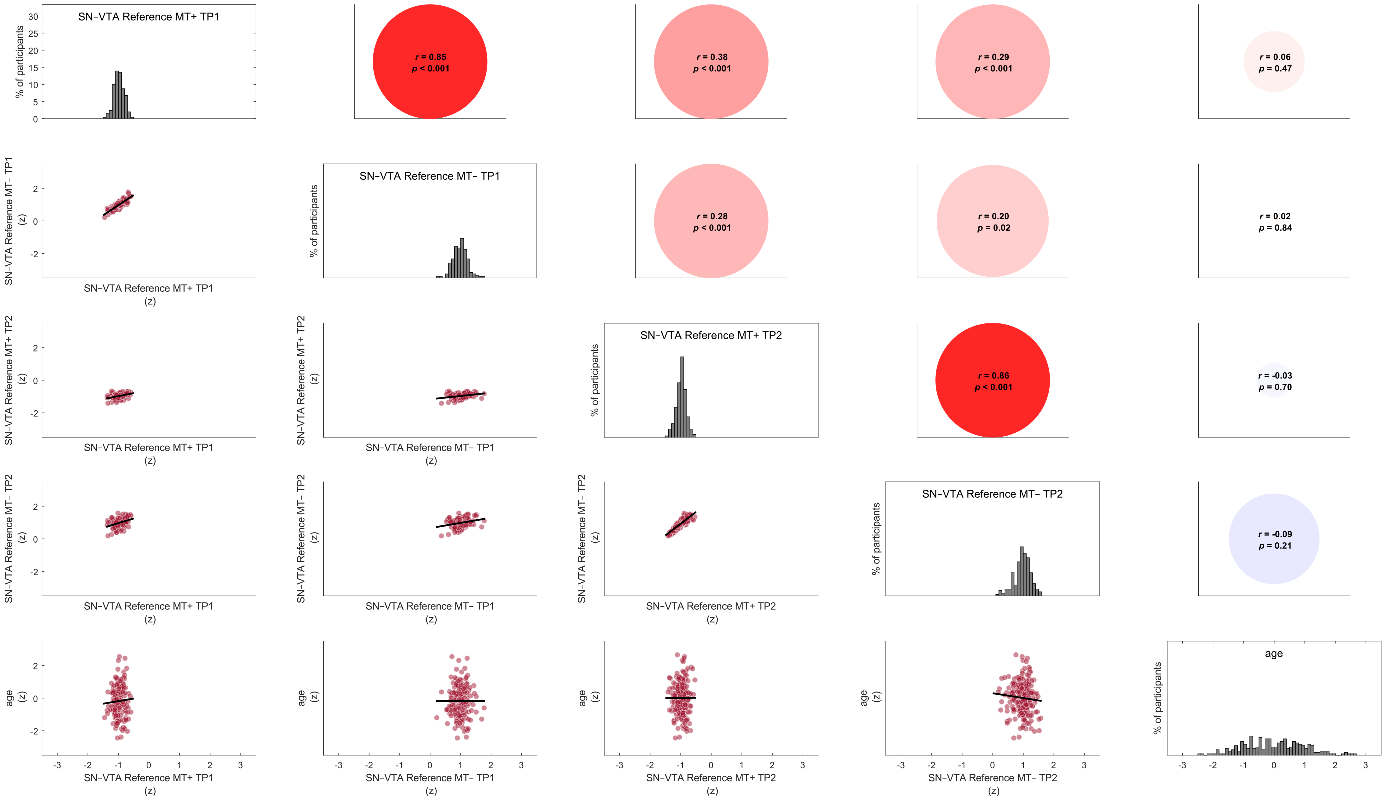

**Figure S24.** Crus cerebri reference intensities across imaging modalities and time points, and their association with chronological age (in older adults). Visualized data are averaged across hemispheres. For the same analyses using SN–VTA data, see Figure S20. Note, the diagonal shows intensity, standardized across all sequences and time points, to facilitate comparing intensity distributions (age was standardized separately). Imaging sequences included a Magnetization Transfer sequence, acquired once with a dedicated magnetic saturation pulse (MT+) and once without, resulting in a proton density image (MT–). LC, locus coeruleus; SN–VTA, substantia nigra–ventral tegmental area; TP, time point.

Longitudinal cognitive models:

**Table S7**. Model fit and invariance for longitudinal cognitive models

| Model number | Model name | Age group | Time point | Invariance | $\chi^2$ | *df* | *p* | RMSEA | CFI |
|---|---|---|---|---|---|---|---|---|---|
| 2.2.1 | Covariance of task-specific WM change factors | OA | 1, 2, 3 | – (single-group, latent-change score models) | 5.005 | 6 | 0.543 | ~ 0 | ~ 1 |
| 2.2.2 | Covariance of task-specific EM change factors | OA | 1, 2, 3 | – (single-group, latent-change score models) | 18.502 | 14 | 0.185 | 0.036 | 0.988 |

*Note:* WM, working memory; EM, episodic memory; YA, younger adults; OA, older adults; RMSEA, root mean square error of approximation; CFI, comparative fit index

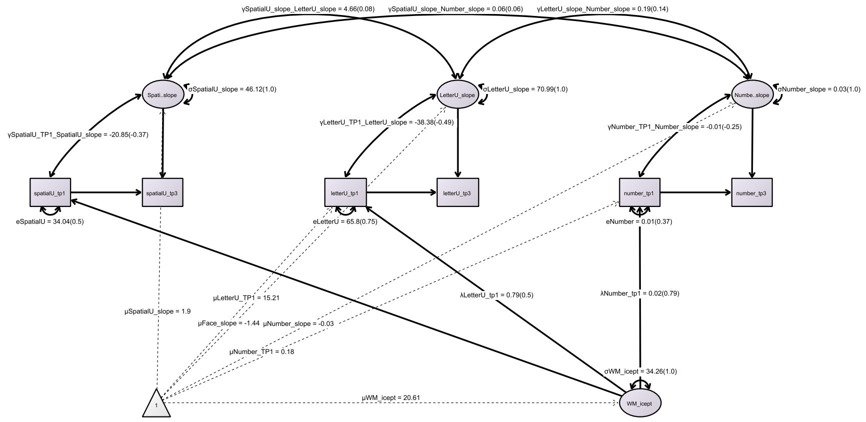

**Figure S25.** Model 2.2.1.

Pictorial rendition of task-specific working memory latent change scores models including data of time point 1 and 3 of older adults. Rectangles and circles indicate manifest (observed) and latent variables, respectively. The constant is depicted by a triangle. (Co)Variances (γ, σ) and loadings (λ) in brackets indicate standardized estimates. One-headed arrows indicate regressions, double-headed arrows indicate correlations. WM, working memory; TP, time point.

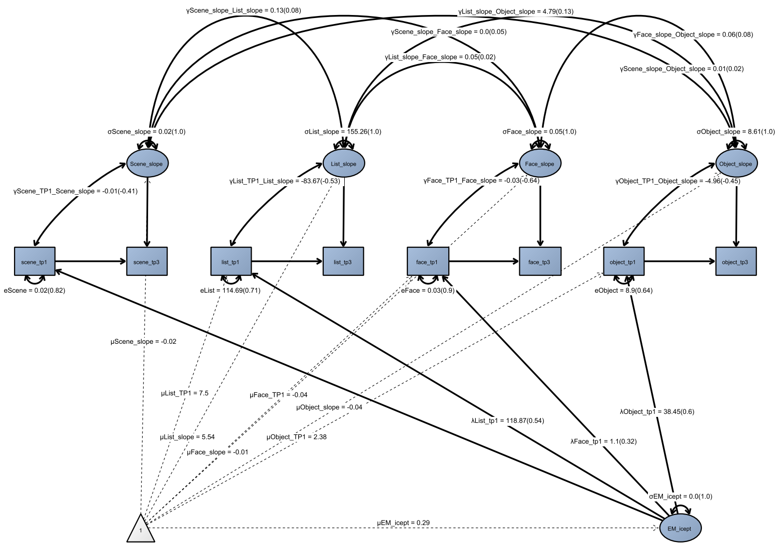

**Figure S26.** Model 2.2.2.

Pictorial rendition of task-specific episodic memory latent change scores models including data of time point 1 and 3 of older adults. Rectangles and circles indicate manifest (observed) and latent variables, respectively. The constant is depicted by a triangle. (Co)Variances (γ, σ) and loadings (λ) in brackets indicate standardized estimates. One-headed arrows indicate regressions, double-headed arrows indicate correlations. EM, episodic memory; TP, time point.

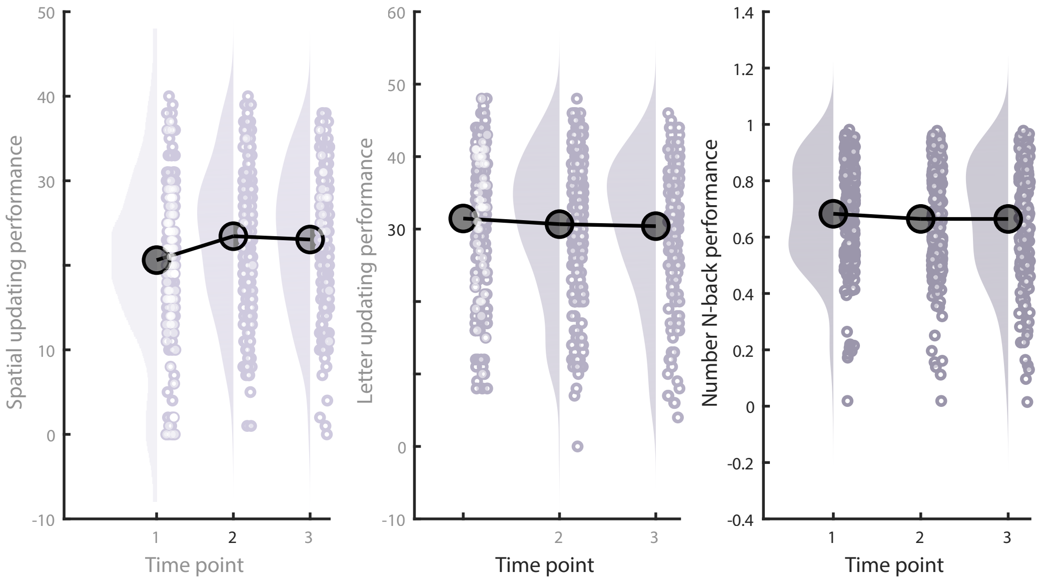

**Figure S27**. Older adults' working memory performance for time points 1–3 for each indicator task. Raincloud plots based on [9].

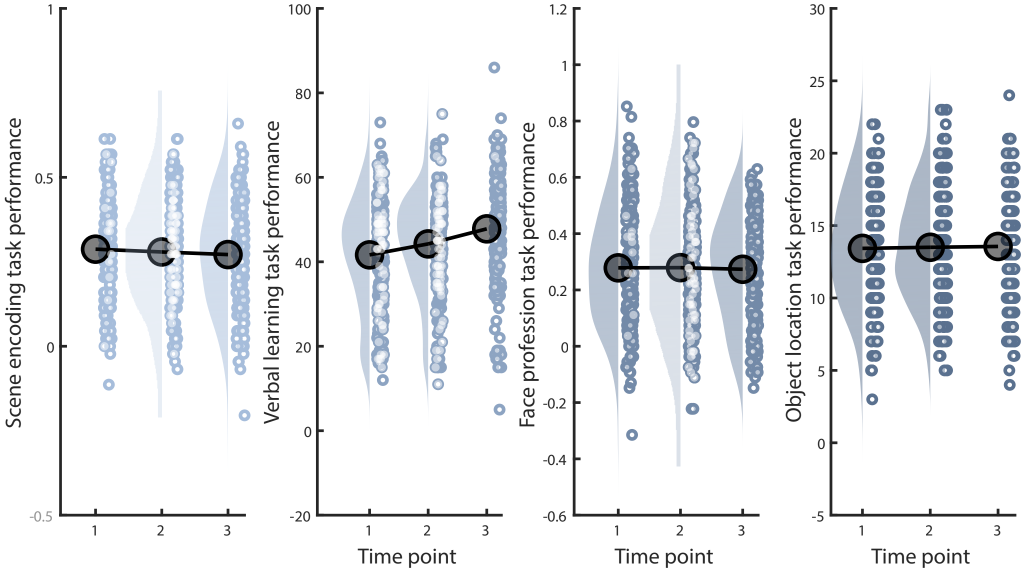

**Figure S28**. Older adults' episodic memory performance for time points 1–3 for each indicator task. Raincloud plots based on [9].

Longitudinal neuro–cognitive models:

**Table S8**. Model fit and invariance for longitudinal neuro–cognitive models

| Model number | Model name | Age group | Time point | Invariance | $\chi^2$ | *df* | *p* | RMSEA | CFI |
|---|---|---|---|---|---|---|---|---|---|
| 2.3.1 | Prediction of EM factor by multimodal LC change factor | OA | 1, 2, 3 | – (single-group, latent-change score model) | 34.799 | 25 | 0.092 | 0.04 | 0.962 |
| 2.3.2 | Prediction of WM factor by multimodal SN–VTA change factor | OA | 1, 2, 3 | – (single-group, latent-change score model) | 20.997 | 18 | 0.28 | 0.026 | 0.984 |

*Note:* LC, locus coeruleus; SN–VTA, substantia nigra–ventral tegmental area; WM, working memory; EM, episodic memory; OA, older adults; RMSEA, root mean square error of approximation; CFI, comparative fit index

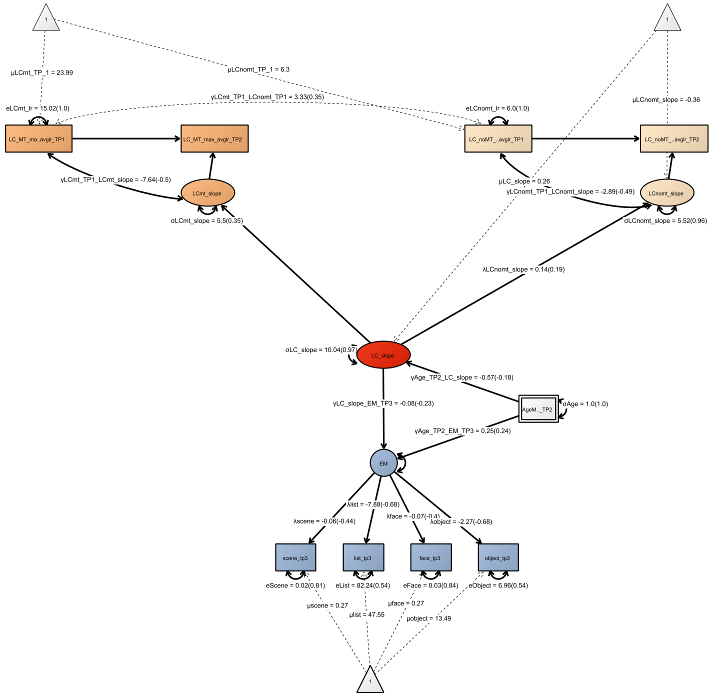

**Figure S29**. Model 2.3.1.

Pictorial rendition of structural equation model predicting episodic memory performance (time point 3) by multi-modal LC change scores (time point 1→2) in older adults.
Rectangles and circles indicate manifest (observed) and latent variables, respectively. The constant is depicted by a triangle. (Co)Variances (γ, σ) and loadings (λ) in brackets indicate standardized estimates. One-headed arrows indicate regressions, double-headed arrows indicate correlations.
LC, locus coeruleus; SN, substantia nigra–ventral tegmental area; fse, Fast Spin Echo; mt, Magnetization Transfer (MT+); nomt, Proton Density (MT–)
EM, episodic memory; TP, time point.

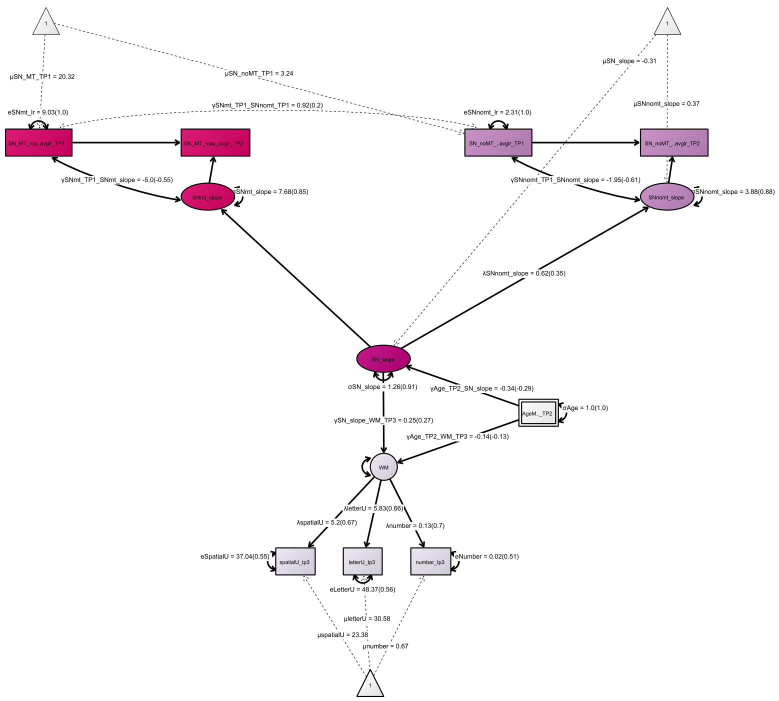

**Figure S30.** Model 2.3.2.

Pictorial rendition of structural equation model predicting working memory performance (time point 3) by multi-modal SN–VTA change scores (time point 1→2) in older adults.

Rectangles and circles indicate manifest (observed) and latent variables, respectively. The constant is depicted by a triangle. (Co)Variances (γ, σ) and loadings (λ) in brackets indicate standardized estimates. One-headed arrows indicate regressions, double-headed arrows indicate correlations.

LC, locus coeruleus; SN, substantia nigra–ventral tegmental area; fse, Fast Spin Echo; mt, Magnetization Transfer (MT+); nomt, Proton Density (MT–); EM, episodic memory; TP, time point.

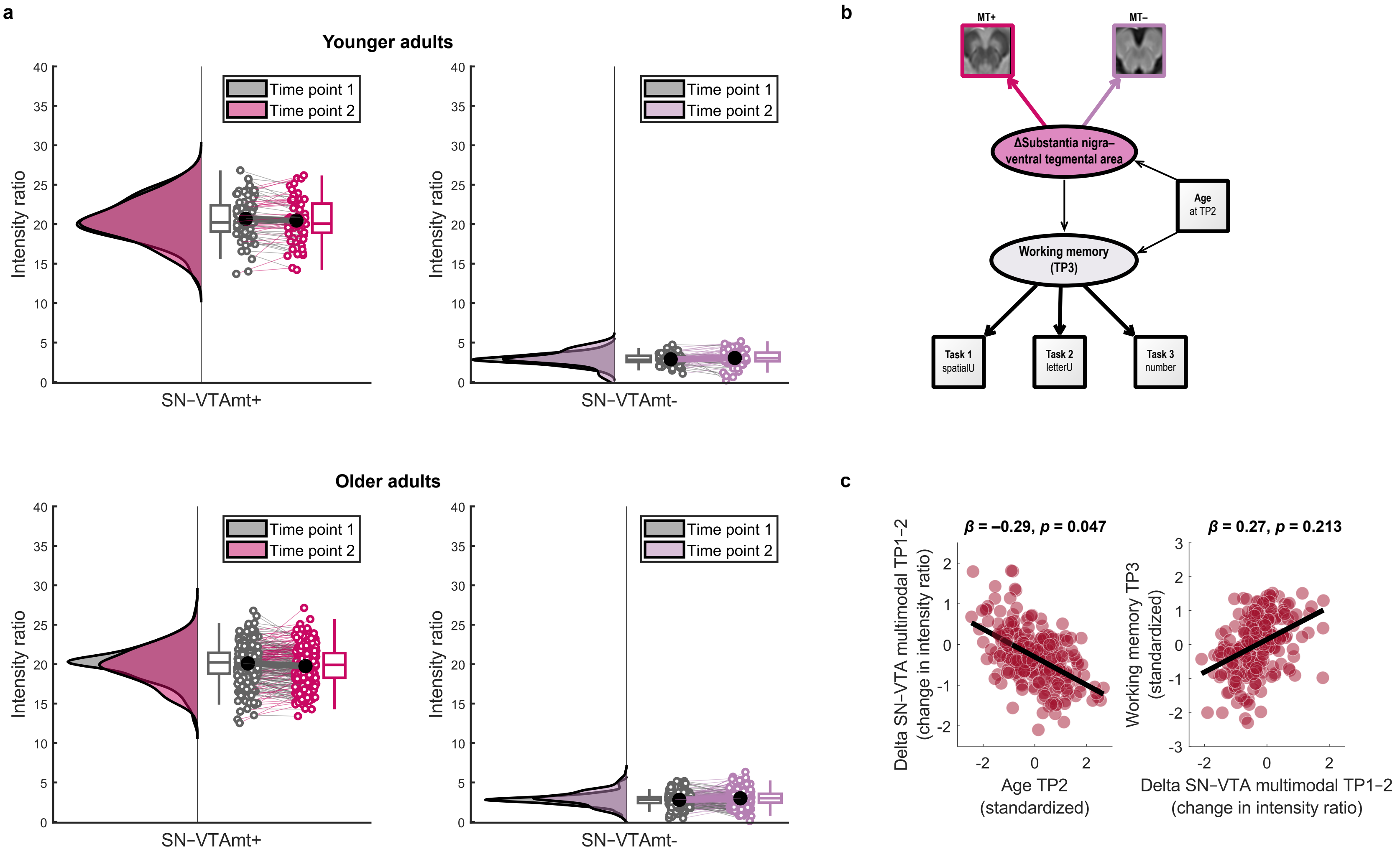
a
Younger adults
Intensity ratio
Time point 1
Time point 2
SN−VTAmt+
SN−VTAmt-
Older adults
b
MT+
MT−
ΔSubstantia nigra–ventral tegmental area
Age at TP2
Working memory (TP3)
Task 1 spatialU
Task 2 letterU
Task 3 number
c
β = −0.29, p = 0.047
β = 0.27, p = 0.213
Delta SN−VTA multimodal TP1−2 (change in intensity ratio)
Age TP2 (standardized)
Working memory TP3 (standardized)
Delta SN−VTA multimodal TP1−2 (change in intensity ratio)

**Figure S31.** Longitudinal changes in SN–VTA intensity ratios and their association with age and future memory performance.

**a**, Numerically, older adults show more negative average change in SN–VTA intensity across time points as compared to younger adults. MRI sequences include a Magnetization Transfer sequence, acquired once with a dedicated magnetic saturation pulse (MT+) and once without, yielding a proton density image (MT–). For the Fast Spin Echo-sequence, only cross-sectional data are available. **b,** Schematic depiction of the structural equation model probing the association of longitudinal change in multimodal SN–VTA integrity with future working memory performance, accounting for chronological age. For the full model, see Figure S24. **c**, Scatter plots showing (1) more negative SN–VTA change in older adults of higher age and (2) the association of future memory performance and SN–VTA change (controlling for chronological age). For comparable analyses using LC and episodic memory data, see Figures S23 and 7. Raincloud plots based on [9]. LC, locus coeruleus; SN–VTA, substantia nigra–ventral tegmental area.

### Cross-sectional and longitudinal neuro–cognitive models with additional covariates:

After establishing relations between catecholaminergic integrity and late-life memory performance, we tested whether these remained significant when accounting for potential confounds. Specifically, we included age and education as standardized covariates in our cross-sectional and longitudinal models (cf. models 1.3.2b, 2.3.1 and 2.3.2) and specified regression paths between these covariates and all neural and cognitive factors.

**Table S8**. Model fit and invariance for neuro–cognitive models with additional covariates

| Model number | Model name | Age group | Time point | Invariance | $\chi^2$ | *df* | *p* | RMSEA | CFI |
|---|---|---|---|---|---|---|---|---|---|
| 1.3.2[1] | Regressions between: MTL, LC, SN–VTA and WM, EM, Gf factors; including age and education as covariates | OA | 2 | – (single-group, cross-sectional model) | 424.121 | 269 | < 0.001 | 0.048 | 0.938 |
| 2.3.1 | Prediction of EM factor by multimodal LC change factor; including age and education as covariates | OA | 1, 2, 3 | – (single-group, latent-change score model) | 45.056 | 33 | 0.079 | 0.038 | 0.954 |
| 2.3.2 | Prediction of WM factor by multimodal SN–VTA change factor; including age and education as covariates | OA | 1, 2, 3 | – (single-group, latent-change score model) | 32.103 | 25 | 0.155 | 0.034 | 0.965 |

*Note:* LC, locus coeruleus; SN–VTA, substantia nigra–ventral tegmental area; WM, working memory; EM, episodic memory; OA, older adults; RMSEA, root mean square error of approximation; CFI, comparative fit index
[1] Model shows Heywood case for σHipp.

We obtained qualitatively similar results to those reported in the main text. That is, cross-sectionally LC integrity was still associated with episodic memory, whereas SN–VTA integrity was related to working memory performance in older adults ($\beta = 0.44$; $\Delta\chi^2(df = 1) = 4.4$; $p < 0.001$ for older adults' LC; $\beta = 0.28$; $\Delta\chi^2(df = 1) = 4.4$; $p = 0.022$ for older adults' SN–VTA). Longitudinally, older adults' LC changes were associated with subsequent episodic memory ($\beta = 0.3$; $\Delta\chi^2(df = 1) = 5.08$; $p = 0.024$ for older adults' LC), whereas the association between SN–VTA change and working memory remained non-significant ($\beta = 0.29$; $\Delta\chi^2(df = 1) = 1.88$; $p = 0.17$ for older adults' SN–VTA).

Finally, to rule out the possibility that unexplored sex effects [11,12] or our treatment of missing values [13] could bias our interpretation, we made use of a different analytical framework (behavioral partial least squares correlation [14–16]) to test for latent brain–behavior associations. These control analyses relied on the same cognitive and neural indicators as our main analyses.

Cross-sectionally, we again found latent associations between the LC and episodic memory ($r = 0.367$; $p < 0.001$) as well as SN–VTA integrity and working memory ($r = 0.218$; $p = 0.004$), which remained significant when including age, education, and sex as covariates ($r_{partial} = 0.256$; $p_{partial} = 0.001$ for older adults' LC; $r_{partial} = 0.199$; $p_{partial} = 0.009$ for older adults' SN–VTA; for comparable statistical approaches, see [16–18]). In addition, longitudinal analyses showed a latent association ($r = 0.345$; $p = 0.036$) between episodic memory performance at time point 3 and changes in LC integrity (conceptualized as difference scores (TP 2–1) for each imaging modality). This association also remained significant when additionally controlling for age, education, and sex ($r_{partial} = 0.325$; $p_{partial} = 0.003$; cf. [17,18]). Taken together, our main and control analyses converge and indicate robust associations between catecholaminergic integrity and memory performance that remain significant when controlling for additional covariates.

### Spatial variation in longitudinal sampling of locus coeruleus and substantia nigra–ventral tegmental area intensity

To test if the position from which intensity values were sampled influenced change analyses, we re-extracted peak intensity ratios for MRI sequences that were assessed at time point 1 and 2 (i.e., MT+, MT–) for each neuromodulatory system, along with their spatial coordinates (x, y, z in MNI space). We then computed the Euclidian distance between the spatial positions from which we sampled at time point 1 and time point 2, using:

$$\text{distance}_{TP1,\,TP2} = \text{sqrt}((TP2_x - TP1_x)^2 + (TP2_y - TP1_y)^2 + (TP2_z - TP1_z)^2)$$

Distance values were then averaged across hemispheres and MRI sequences (MT+, MT–) and compared across neuromodulatory systems. Importantly, we did not find evidence for a higher spatial deviance for the SN–VTA as compared to the LC (Wilcoxon signed rank test; $Z = –0.641$, $p = 0.521$; see below). For the majority of participants, intensity values were extracted from highly comparable locations across time points (distance $\leq$ 3 mm, which may correspond to one voxel (native resolution: 1 × 1 × 3 mm).
At this point, we would like to emphasize that the distance measure reported here has a different meaning than distance measures commonly used in functional MRI analyses, such as, frame-wise displacement. That is, a homogenous hyperintensity distribution within the LC and SN–VTA search spaces could lead to sampling from different spatial positions over time even without movement (cf. Figure 3 in the main text for a visualization of the hyperintensity on a sample level; the corresponding MRI templates are available via [19]). To support this argument, we tested whether sampling from different spatial positions over time would be related to changes in intensity estimates (as could be assumed for movement in the scanner). Neither for the LC nor for the SN–VTA we found an association between Euclidian distance and changes in intensity estimates ($p$s > 0.19; see below).

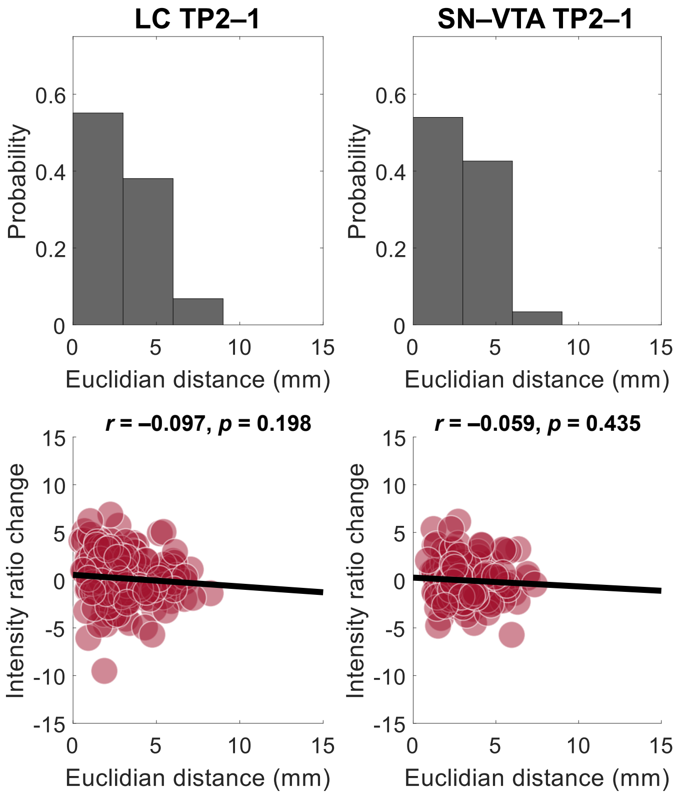

**Figure S32.** Euclidian distance of spatial positions from which intensity ratios were sampled at time point 1 and 2 for the locus coeruleus and substantia nigra–ventral tegmental area.

Euclidian distance did not differ significantly across neuromodulatory systems (Wilcoxon signed rank test; Z = –0.641, p = 0.521) and was not associated with intensity changes for either neuromodulatory system ($p > 0.19$).
